# Structures of LolB bound to LolA or lipoprotein provide insight into the final steps of bacterial lipoprotein trafficking

**DOI:** 10.64898/2026.09.01.748475

**Authors:** Abigail E. Jepson, Elise Kaplan, Nicholas P. Greene, Vassilis Koronakis

## Abstract

In Gram-negative bacteria, lipoproteins are structural elements of the outer membrane and essential components of machineries responsible for its construction and maintenance. The Lol system, responsible for the trafficking of lipoproteins from the site of maturation on the inner membrane to the outer membrane, is therefore crucial to the function of the cell envelope and a key target of efforts to find novel antimicrobials. In the final steps of this process, the outer membrane receptor, LolB accepts triacylated lipoproteins from the periplasmic chaperone LolA before inserting them into the outer membrane. Here we present a structure of LolB in complex with LolA, validated by *in vivo* and *in vitro* assays, highlighting how positively charged residues on the convex face of the LolB β-barrel underpin complex formation. A protruding loop of LolB, essential for function, inserts into the LolA cavity in position to initiate the displacement of substrate lipoprotein from LolA to enable transfer to LolB. Structural resolution of a lipoprotein-bound LolB complex in combination with biophysical assays shows how a molecular latch releases the lid of the cavity to accommodate the lipoprotein acyl chains. Modelling of these structures onto computationally predicted orientations for LolB on the outer membrane provides a rationale for LolA release and lipoprotein triacyl group membrane insertion. Taken altogether, our data elucidate atomic resolution of two key intermediates and provide a greater understanding of the terminal steps of lipoprotein trafficking events at the bacterial outer membrane.

## Introduction

The outer membrane (OM) of Gram-negative bacteria is a selectively permeable barrier vital to the function of the cell. This asymmetric bilayer comprises inner-leaflet phospholipids and outer-leaflet lipopolysaccharide (LPS) that restrict entry of noxious compounds including antibiotics, while protein components allow the import of essential nutrients. Maintenance of this barrier is underpinned by the Lpt, Bam, and Mla machineries, respectively responsible for insertion of LPS (1), β-barrel proteins (2), and maintenance of phospholipid asymmetry (3). These systems are dependent on OM lipoproteins, a class of soluble proteins tethered to the membrane by an N-terminal triacyl anchor (4–6). Lipoproteins also perform roles in maintenance of the structural stability of the cell envelope (7, 8), nutrient acquisition (9), host-pathogen interactions (10), detoxification of antibiotics (11) and response to stress (12). Taken altogether, lipoproteins are essential for bacterial cell physiology and interfering with their synthesis or localization results in cell death (13, 14).

Lipoproteins are synthesized in the cytosol and targeted to the inner membrane (IM) by an N-terminal signal peptide that facilitates translocation by the Sec or Tat pathways (15, 16) and contains a four-residue lipobox motif which directs triacylation of an invariant cysteine at the fourth position of the motif. Sequential action of Lgt, Lsp and Lnt enzymes causes addition of a diacyl group to the cysteine thiol, removal of the signal sequence, and acylation of the amino terminus resulting in the mature lipoprotein. Most lipoproteins are directed to the OM by a dedicated transport pathway, the Lol system (17) except those carrying a transport avoidance motif, commonly aspartate in *E. coli*, immediately following the triacylated cysteine (18). In *E. coli*, the Lol apparatus is composed of the IM ATP-binding cassette (ABC) transporter LolCDE, the periplasmic chaperone LolA and the OM receptor LolB, itself a lipoprotein (19–21). Genetic manipulation of *E. coli* revealed an alternative LolA-LolB independent pathway (22) but under normal conditions all the Lol proteins are essential for cell viability (19–21). The Lol pathway is therefore a key target for the development of novel antibacterials (23–26).

Recent structural studies have elucidated details of discrete events within the transport cycle (Figure 1). LolCDE recruits LolA via interactions with a β-hairpin ‘Hook’ and three surface residue ‘Pad’ in the LolC periplasmic domain (27). Lipoproteins are recognized by LolCDE via their triacyl chains, which are accommodated at the interface of the LolC and LolE subunits (28, 29). ATP binding and subsequent hydrolysis induces the closure of the transporter by a mechanotransmission mechanism typical of type VII ABC transporters (30, 31), pushing the triacyl moiety towards LolA. The LolA-lipoprotein complex, in which the acyl chains are accommodated inside the LolA cavity (32), is then released to the periplasm before the LolCDE transporter resets to an open state. In the final step, the OM receptor LolB accepts the lipoprotein from LolA and inserts it into the phospholipid layer (19, 33).

**Figure 1.**
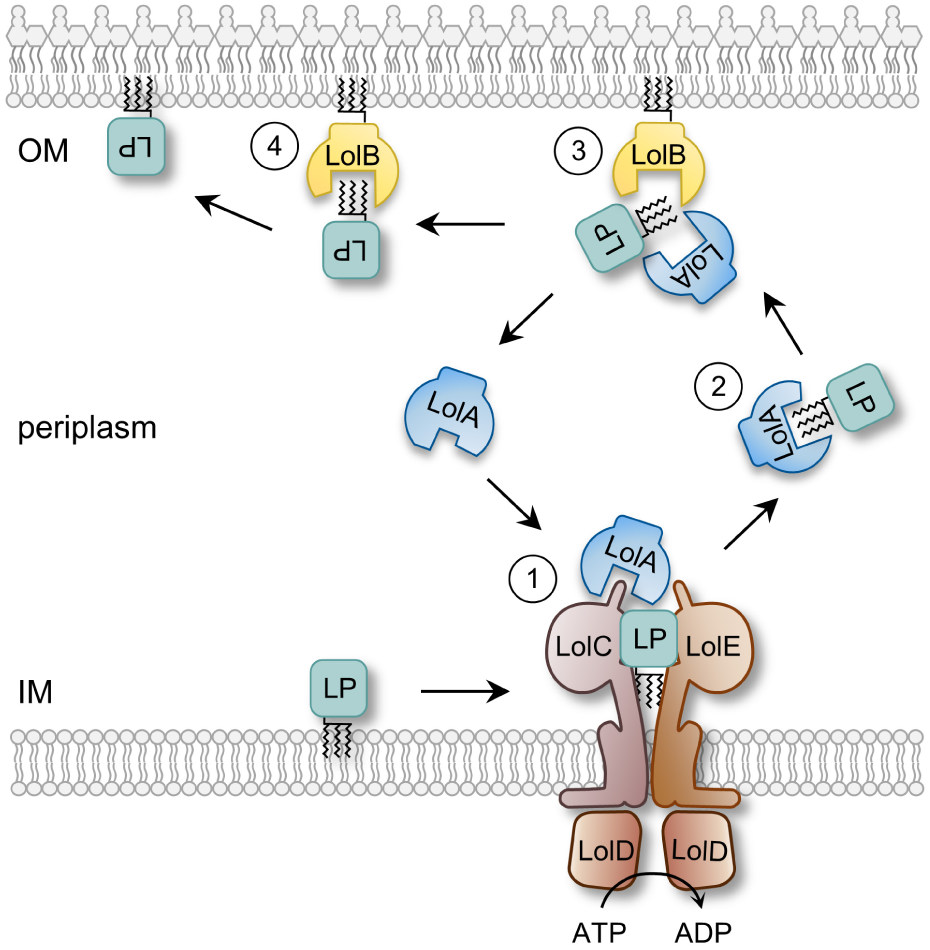
Lipoprotein trafficking in *E. coli* via the Lol pathway. 1) LolA is recruited by the LolCDE ABC transporter through interaction with the ‘Hook’ and ‘Pad’ structural features of LolC (27). Lipoprotein (LP) acyl chains bind at the interface of LolC and LolE (28). Binding of ATP to the nucleotide-binding protein LolD and subsequent nucleotide hydrolysis triggers lipoprotein transfer to LolA and resets the LolCDE transporter for a new cycle. 2) The lipoprotein acyl chains are enclosed inside the LolA cavity (32) forming a soluble complex able to cross the periplasm. 3) The lipoprotein-LolA complex interacts with outer membrane anchored-LolB, and lipoprotein transfer occurs between the LolA and LolB proteins. 4) The hydrophobic cavity of LolB is hypothesized to accommodate the lipoprotein acyl chains prior to insertion into the inner leaflet of the outer membrane.

LolA and LolB both comprise a half β-barrel with three α-helices that form a lid over a hydrophobic central cavity (34). Protein-specific roles are directed through distinct features; the extended C-terminus of LolA, absent from LolB, is essential for interaction with LolC (27, 35) whereas LolB possesses a protruding loop, tipped by the L68 residue (34) that is required for both acceptance and membrane insertion of lipoproteins (36). The apparent absence of LolB in some organisms is compensated by the presence of a protruding loop in LolA rendering the protein capable of performing both functions (37). Lipoprotein transfer from LolA to LolB occurs in a unidirectional manner independently of energy input (38). Based on NMR and *in vivo* crosslinking experiments, a ‘mouth-to-mouth’ orientation has been proposed where the two protein cavities align in a manner that could allow transfer of the lipoprotein acyl chains (39, 40).

Here we present the crystal structure of LolB in complex with LolA, validated by biochemical and *in vivo* data. By exploiting a lipoprotein transfer assay, we isolated a lipoprotein-bound LolB complex revealing how the LolB cavity opens to accept lipoprotein substrate from LolA. In combination with existing computational data, we propose a model for how LolB receives lipoprotein from LolA and facilitates its integration into the membrane. Our data reveal molecular details of two discrete events in the lipoprotein trafficking pathway at the outer membrane and may provide new targets for antimicrobial therapy.

## Results

### Crystal structure of *E. coli* LolA R43L-LolB complex

Deciphering the interaction between LolA and LolB is crucial to better understand lipoprotein transfer. LolB displays a 10-fold greater affinity for LolA R43L, a mutant that mimics a lipoprotein-bound state, than for the wild-type protein (32), making it a more suitable candidate for creation of stable complexes. We therefore set up crystallization trials with soluble, non-acylated LolB and LolA R43L that was either in the free-state or complexed with lipoprotein. We were not able to crystallize the LolA-lipoprotein-LolB ternary complex but we obtained a crystal of the LolA R43L-LolB complex in space group P*22_1_2_1_* that diffracted to a resolution of 2.37 Å. The overall structure of the complex is shown in Figure 2A with X-ray data and refinement statistics listed in *SI Appendix* Table S1. Movie S1 shows a rolling tour of the complex along with representative electron density.

**Figure 2.**
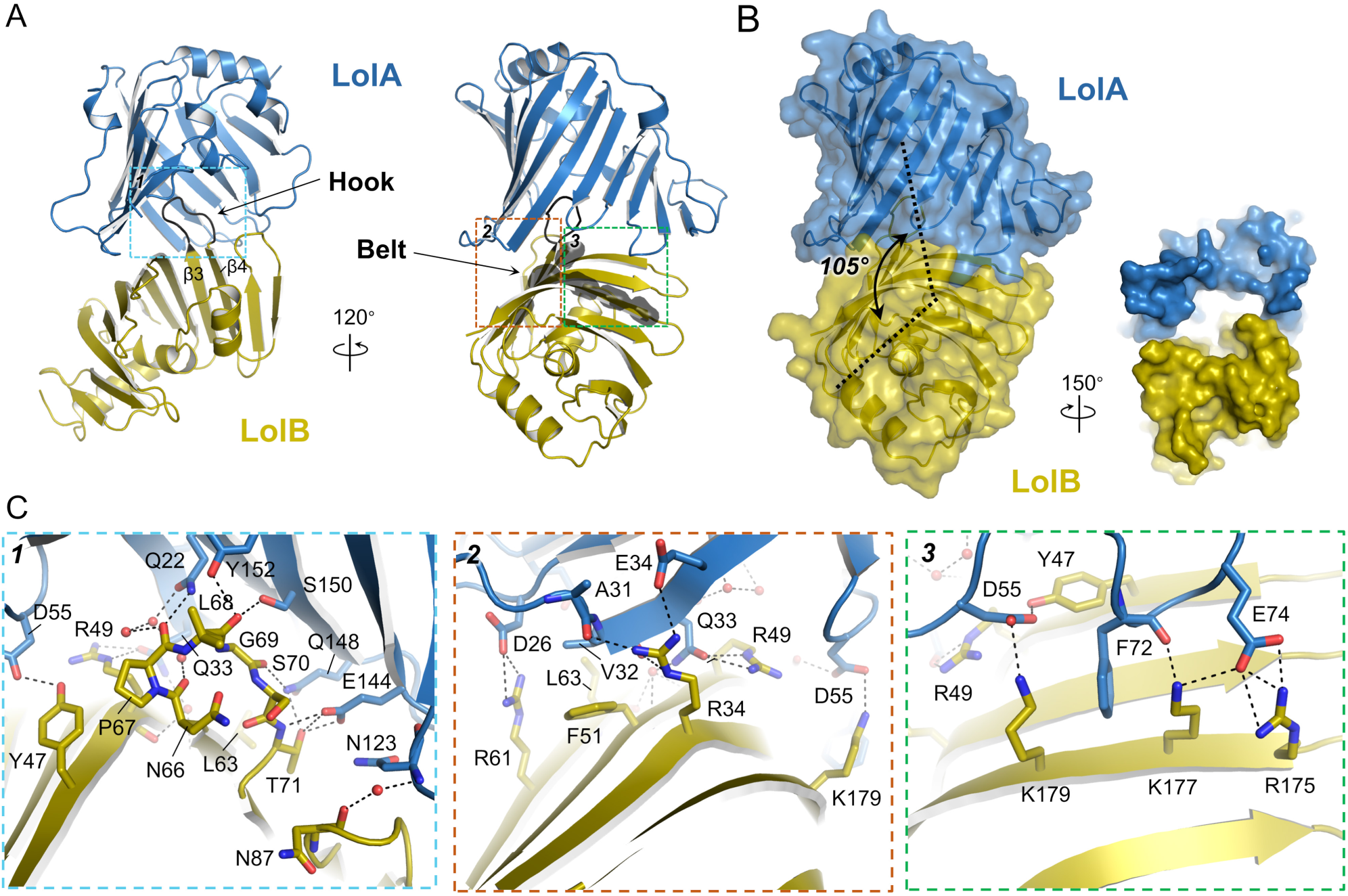
Crystal structure of LolB bound to LolA R43L. (A) Overall views of the complex with LolA R43L in blue and LolB in yellow. The positions of LolB ‘Hook’ and ‘Belt’ are indicated. Main chains of LolB Belt residues are shown as black transparent surface. (B) Surface representation of the LolA R43L-LolB complex. Dashed lines indicate the axis of each cavity, highlighting the 105° angle between the two protein barrels. A view of the LolA-LolB cavity entrance is shown in inset. (C) Close-up views of the boxed regions in A showing the interactions between LolA and LolB.

The β-barrels of LolA R43L and LolB are aligned with the two cavities facing each other, consistent with the mouth-to-mouth model previously described by Okuda *et al.* (40) but with the axis of each cavity meeting at an angle of 105° (Figure 2B). This arrangement creates a continuous chamber that extends across the two protein cavities to enable transfer of the lipoprotein acyl chains while leaving sufficient space for the lipoprotein peptide chain to extend out (Figure 2B, *inset*). Both LolA and LolB within the complex adopt a closed conformation, aligning with a rmsd of 1.4 Å over 177 C_α t_o closed LolA R43L (PDB 2ZPC, (41)), and 0.9 Å over 176 C_α t_o wild-type LolB (PDB 1IWM, (34)), respectively.

The complex is stabilized by multiple interactions between residues located at the concave face of LolA R43L and at the convex surface of LolB as shown in Figure 2A, C. About 12% of each protein surface is buried in the complex interface. The interaction is supported by the insertion of a LolB protruding loop (residues T65-T71), which links the β3 and β4 strands, into the cavity of LolA and via a number of positively charged residues located on the LolB convex face. This mode of binding is highly reminiscent of LolA recruitment by a β-hairpin loop (the ‘Hook’) and a trio of surface residues (the ‘Pad’) in LolC (27). By analogy with LolC, we refer to the hairpin loop as the LolB ‘Hook’. The LolC ‘Pad’ includes two arginine residues while here, LolB-LolA interaction is further mediated by multiple basic residues (R34, R49, R61, R175, K177 and K179) located at the LolB surface. We define this series of charged residues as the LolB ‘Belt’ as they span the entire interface and interact with three pairs of β-strands in LolA.

The LolB Hook is stabilized inside the LolA cavity by van der Waals (VDW) interactions mediated by P67 and L68 at the tip of the Hook and by a series of hydrogen bonds engaging the main chains of all the Hook residues but S70 (Figure 2C, panel 1). Additionally, the side chain of T71 interacts with LolA E144 and Q148. The LolB Belt brackets LolA by means of five salt bridges. On one side of the LolB convex face, R34 and R61 engage with LolA E34 and D26 respectively (Figure 2C, panel 2), while on the other side the R175/K177/K179 triad interacts with LolA E74 and D55 residues via three additional salt bridges (Figure 2C, panel 3). A cation-pi sandwich is formed between the side chains of LolB K177 and K179 and the aromatic ring of LolA F72. The sixth Belt residue, R49, contributes to the complex stability via hydrogen bonds with LolA Q33.

We mapped previously reported *in vivo* photo-crosslinking data (40) onto our structure revealing good agreement (∼ 94%) between the two approaches. Residues in LolA or LolB which form photo-inducible crosslinks when replaced by the unnatural amino acid benzoyl-phenylalanine, are clustered at the interface between the two proteins (*SI Appendix* Figure 1A, red), while residues distant from the interaction do not form crosslinks (*SI Appendix* Figure 1A, blue). The structure is comparable to that of a recently published wild-type LolA-LolB complex from *Xanthomonas campestris* (42). In both the *X. campestris* and *E. coli* structures, the relative orientations of LolA and LolB are similar (*SI Appendix* Figure 1B) and the two complexes can be aligned with an rmsd of 2.5 Å over 314 C_α_, suggesting that the R43L mutation does not affect the complex structure. The *X. campestris* complex is also stabilized via interaction of LolA with the LolB Hook and charged residues analogous to the *E. coli* LolB Belt (*SI Appendix* Figure 1C). Residues K186 and R188 form interactions analogous to R175 and K177 in *E. coli* with additional interactions provided by R45 and R177 from adjacent β-strands. Fewer interactions are present on the other side of the LolB Belt in *X. campestris* (*SI Appendix* Figure 1C, D) where the LolA loop joining β5-β6 is stabilized by a single LolB residue, R58.

Residues stabilizing the LolA-LolB structure are primarily situated on the external face of the LolA β-barrel that, unlike the lid helices, does not change in conformation upon lipoprotein insertion (32). The interactions described here would therefore be compatible with the formation of a lipoprotein-bound LolA-LolB complex. In concert with the *in vivo* crosslinking data, this suggests that our structure constitutes an appropriate model for physiological interactions between LolB and wild-type LolA.

### Mutating residues in the LolB Belt and Hook affects LolA-LolB association *in vitro* and lipoprotein transfer *in vivo*

The LolA R43L-LolB complex is stabilized by hydrogen bonds and salt bridge interactions involving the side chains of eight LolB residues (Figure 2C). To establish the importance of each residue in the complex formation, we produced individual alanine variants and first evaluated their ability to associate with wild-type LolA, and then to accept the lipoprotein Pal from LolA. Results are presented in Figure 3A with raw data for the association and transfer assays provided in *SI Appendix* Figure 2. All LolB mutants exhibited a reduced ability to bind LolA relative to wild-type LolB. Alanine substitution of residue K177 resulted in the most significant effect, reducing LolA binding by more than 85%, consistent with its interaction with two residues in LolA. However, none of the mutations significantly reduced the ability of LolB to accept lipoprotein from LolA, likely because transient interactions permit lipoprotein transfer.

**Figure 3.**
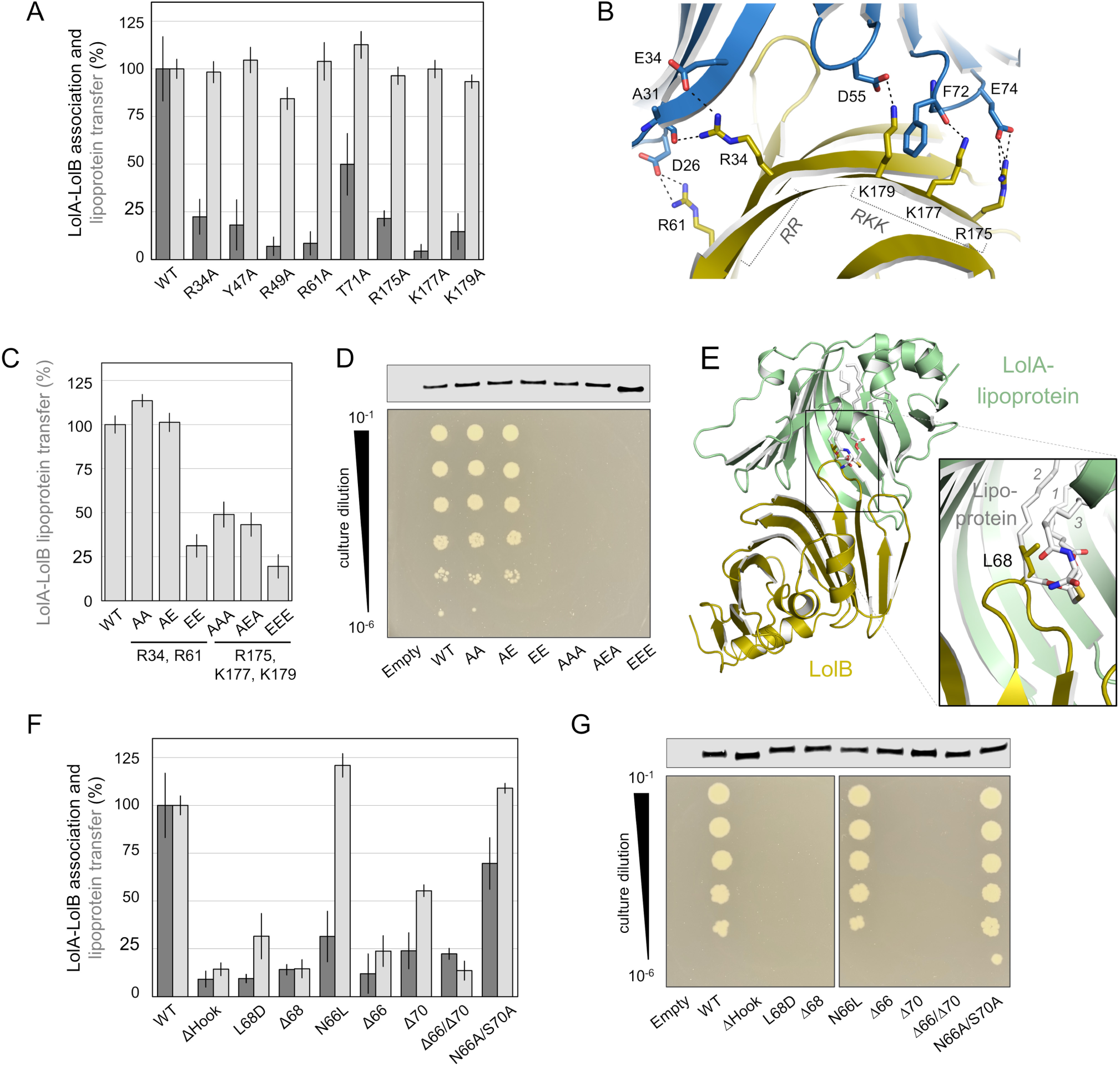
Role of the LolB Hook and Belt in the LolA-LolB interaction. (A) *In vitro* interaction and lipoprotein transfer (dark and light grey respectively) between LolA and wild-type LolB (WT) or indicated variants. His-tagged LolB proteins were incubated with tag-free LolA or LolA-Pal complex prior to immobilization on Ni resin. After washing, bound proteins were eluted and quantified by SDS-PAGE. Data were normalized against the value obtained for wild-type LolB and shown as the mean ± standard deviation for triplicate experiments. (B) Close-up view of LolA R43L-LolB structure highlighting the two clusters of charged residues R34 and R61 (RR) and R175, K177 and K179 (RKK) on the convex side of the LolB β-barrel. (C) Transfer of Pal lipoprotein from LolA to LolB was assessed as described in (A) for wild-type LolB or variants containing alanine or glutamate mutations in the two charged clusters RR and RKK. (D) Expression of plasmid-borne wild-type *lolB* or variants in BW65 cells supported by expression of chromosomal *lolB* (top). Serial dilutions of a conditional *lolB* knockout *E. coli* BW65 strain carrying either plasmid-borne wild-type *lolB* or indicated variants in the absence of inducer required for expression of chromosomal *lolB* (bottom). (E) Structural alignment of lipoprotein-bound LolA (7Z6W) with the LolA R43L-LolB complex highlighting the steric clash of LolB Hook with the lipoprotein ligand. Acyl chain numbering is indicated. (F) Association with LolA and transfer of Pal lipoprotein from LolA to LolB as described in (A) for wild-type LolB or variants in the Hook. (G) Expression of LolB Hook mutants and their ability to support cell growth in the absence of chromosomal *lolB* evaluated as in (D). Controls for panels D and G demonstrating growth in the presence of chromosomally-encoded wild-type *lolB* can be found in *SI Appendix* Figure 3.

The LolB Belt comprises two charged clusters, consisting of R34/R61 and R175/K177/K179, hereafter RR and RKK respectively (Figure 3B), raising the possibility that redundancy within these clusters could mask effects of single mutations. We therefore engineered three double mutants in the RR cluster (AA, AE and EE) and three triple mutants in the RKK cluster (AAA, AEA and EEE) by replacing the basic residues with alanine and/or glutamate. The ability of these variants to accept Pal from LolA was then assessed (Figure 3C). In the RR cluster, only reversal of both charges (EE) had a significant effect on lipoprotein transfer. The RKK cluster was more sensitive to mutation; triple AAA and AEA mutants only retained 50% activity while lipoprotein transfer decreased by almost 80% for the EEE variant. In *E. coli, lolB* is essential (43), so to probe the importance of these charged clusters *in vivo* we examined the ability of the double and triple mutants to support growth in a *lolB* conditional knockout strain. Consistent with the *in vitro* results, the AA and AE variants were able to support growth in the absence of wild-type *lolB* but the EE and all the mutants in the RKK cluster could not (Figure 3D). Overall, our *in vitro* lipoprotein transfer and *in vivo* results demonstrate that the RKK cluster, in particular, has an important role in the physiological function of LolB.

Alignment of lipoprotein-bound LolA with our LolA-LolB complex shows that the Hook apex residue, L68, sterically clashes with the lipoprotein R2 acyl chain inside the LolA cavity (Figure 3E). This clash suggests that the Hook initiates the transfer by physically dislodging the lipoprotein from LolA. We created mutations in the Hook and assessed the ability of the variants to interact with LolA, receive lipoprotein from LolA and support bacterial growth (Figure 3F, G). Removal of the entire Hook (ΔHook) or deletion (Δ68) or glutamate substitution of the tip residue, L68, severely impacted *in vitro* lipoprotein transfer and association with LolA, consistent with the inability of these variants to sustain cell growth in the absence of wild-type *lolB*.

The Hook is stabilized by a hydrogen bond between the side chains of LolB residues N66 and S70 (Figure 2C). Abrogating this interaction by substitution of N66 with leucine reduced association with LolA by 50% but neither lipoprotein transfer nor bacterial growth were affected. In contrast, removal of N66 (Δ66) was lethal and we reasoned that the consequent displacement of L68 was likely responsible for the abrogation of function. Consistent with this hypothesis, deletion of the Hook residue opposite N66, S70 (Δ70) or concomitant removal of residues N66 and S70 (Δ66/Δ70) disrupted LolA association, lipoprotein transfer and consequently the ability to maintain growth. Conversely, maintaining the Hook length and structure while removing the hydrogen bond between N66 and S70 side chains (mutant N66A/S70A) permits lipoprotein transfer and fully supports growth. Taken altogether our structural, biochemical and *in vivo* data demonstrate that the Hook provides a precise scaffold to project a hydrophobic residue into the LolA cavity where it would be in position to displace the substrate triacyl chains to initiate transfer to LolB.

### A molecular latch controls access to the LolB cavity

In the structure described here and that previously published (34), LolB adopts a closed conformation with the helical lid occluding the protein cavity. To understand how the opening of the cavity is controlled, we identified residues that stabilized the helical lid in a closed position through either hydrogen bonds (Q44, Q57, E98, Q116, Q143) or hydrophobic interactions (L114, I118) with residues of the β-barrel (Figure 4A). We then abrogated these interactions by mutation to alanine or glycine as appropriate and assessed the cavity state of the LolB variants by utilizing the fluorescent fatty acid probe, 11-(Dansylamino) undecanoic acid (DAUDA), that exhibits increased and blue shifted fluorescence emission upon binding to a hydrophobic surface (44).

**Figure 4.**
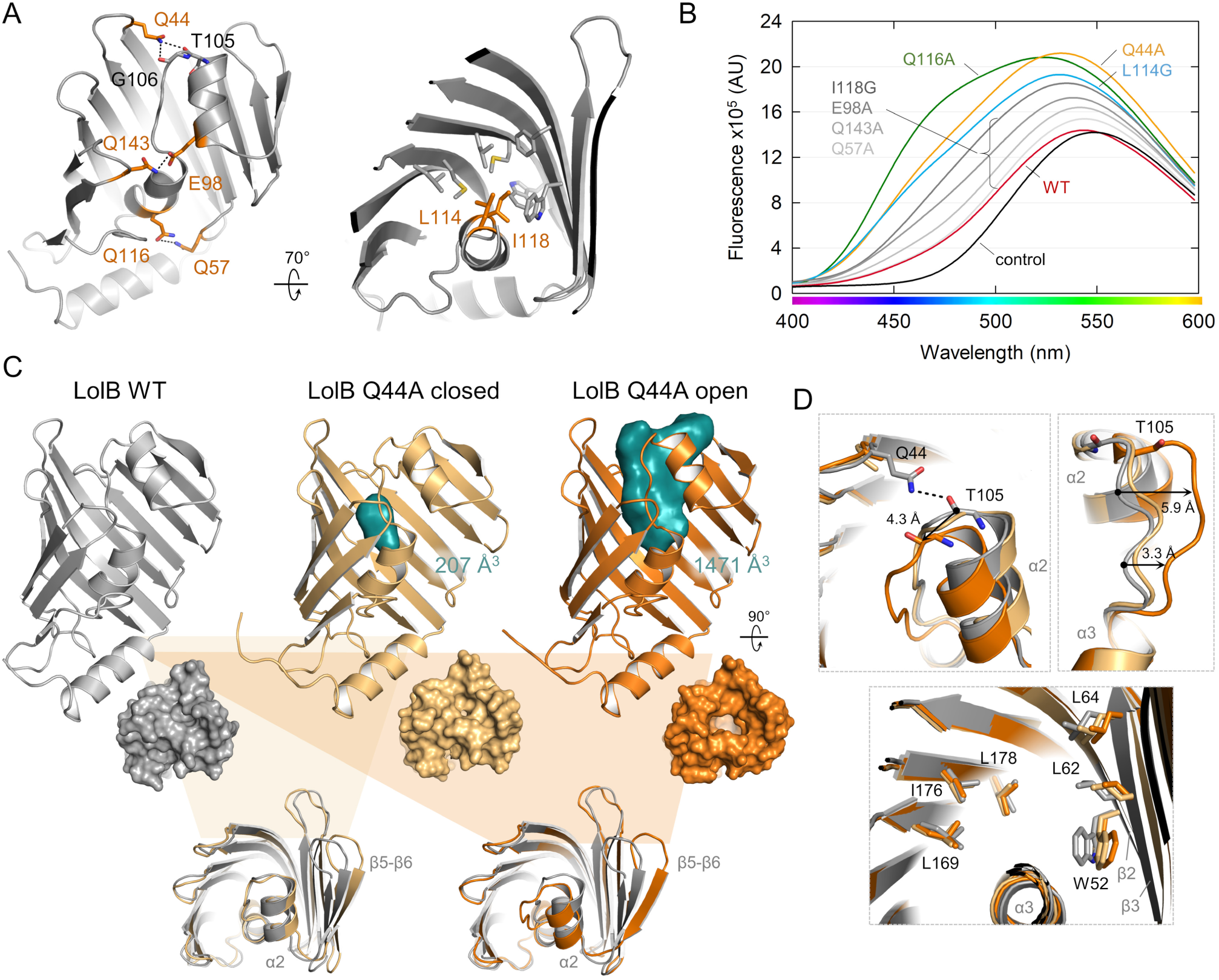
Key residues that control LolB helical lid release. (A) Residues targeted for mutagenesis are shown as orange sticks in wild-type (WT) LolB (grey, 1IWM). (B) Binding of DAUDA to LolB wild-type and variants. The control corresponds to basal DAUDA fluorescence in the absence of protein. (C) Comparison of the structures of LolB WT (grey), and the Q44A mutant in the closed (light orange) and open (dark orange) conformation. Protein cavities are shown as solid teal surfaces. (D) Close-up views of the α2 helix (top left), the α2-α3 linker (top right) and the LolB central cavity (bottom).

Incubation of DAUDA with soluble, wild-type LolB resulted in only a slight increase in fluorescence compared to the buffer control (Figure 4B). A similarly modest increase was observed for variants Q57A, E98A, I118G, Q143, whereas addition of DAUDA to L114G, Q44A and Q116A mutants resulted in greater fluorescence intensities accompanied by a blue-shift. To investigate the structural changes enabling an enhanced interaction with the probe, we crystallized the two variants, Q44A and Q116A, that exhibited the most significant DAUDA binding. X-ray data for multiple LolB Q116A crystals revealed a protein in a closed conformation akin to the wild-type LolB. Conversely, the Q44A mutant crystallized in two distinct conformations: a closed form, solved at 1.88 Å resolution, and an open form, with a larger cavity, diffracting to 2.12 Å. The structures are shown in Figure 4C with X-ray data and refinement statistics listed in *SI Appendix* Table 1.

The wild-type LolB and closed Q44A structures are highly similar, except that we detected a small internal cavity for the LolB Q44A closed conformation. In contrast, the open conformation of LolB Q44A exhibits a surface-accessible cavity with an estimated volume of 1471 Å^3^, a substantial increase compared to the closed conformation. Abrogation of the Q44-T105 hydrogen bond allows ∼5 Å shift of the α2 helix and α2-α3 flexible linker alongside minor movements of the cavity-facing side chains on β2-β3 strands, resulting in a larger cavity (Figure 4D). Taken altogether these results suggest that the Q44-T105 hydrogen bond acts as a ‘latch’ that maintains the protein cavity in a closed conformation in the absence of lipoprotein.

### Crystal structure of the LolB L114G - lipoprotein complex

To understand how the substrate acyl chains are accommodated, we next aimed to crystallize LolB in complex with Pal lipoprotein. However, because LolB spontaneously inserts lipoproteins into lipid membranes, the production of the complex *in vivo* was precluded. We therefore generated lipoprotein-bound LolB *in vitro*, using LolA-lipoprotein complexes produced *in vivo*. Production of sufficient quantities of stable wild-type LolB-lipoprotein complex was challenging. However, we noticed that LolA transferred lipoprotein more efficiently to LolB L114G, resulting in a greater yield of lipoprotein-bound LolB complex (*SI Appendix* Figure 4A). This mutant was still able to support growth at a level comparable to the wild-type protein (*SI Appendix* Figure 4B). Cleavage of the flexible linker and mature domain of Pal as performed for the LolA-lipoprotein complex crystallization (32) enabled crystallization of the LolB L114G-lipoprotein complex.

We determined the crystal structure of soluble LolB L114G in complex with lipoprotein at 1.75 Å. The complex crystallized in space group *P*2_1_2_1_2_1_ with two copies of the LolB-lipoprotein complex per asymmetric unit. In both copies, electron density was resolved for the triacylated cysteine but not for the rest of the peptide chain, suggesting that LolB does not make specific interactions with the lipoprotein globular domain. Movie S2 shows a rolling tour of the LolB-lipoprotein complex for each chain along with representative electron density. The structure is shown in Figure 5A with X-ray data and refinement statistics listed in *SI Appendix* Table 1 and omit maps for the triacyl moiety shown in *SI Appendix* Figure 5. In both copies, the lipoprotein acyl chains are similarly positioned within the hydrophobic LolB cavity except for the final six carbons of R1 that adopt different conformations, possibly reflecting conformational freedom in the absence of the leucine side chain in the L114G mutant (*SI Appendix* Figure 6A). Details of the lipoprotein binding mode are shown in Figure 5B. Residue Q44 forms a hydrogen bond with the R2 ester linkage, while the acyl chains are stabilized by VDW interactions involving multiple residues within the LolB cavity. In comparison with the closed form of LolB, the cavity volume reaches 1982 Å^3^, a value similar to that of lipoprotein-bound LolA (Figure 5C).

**Figure 5.**
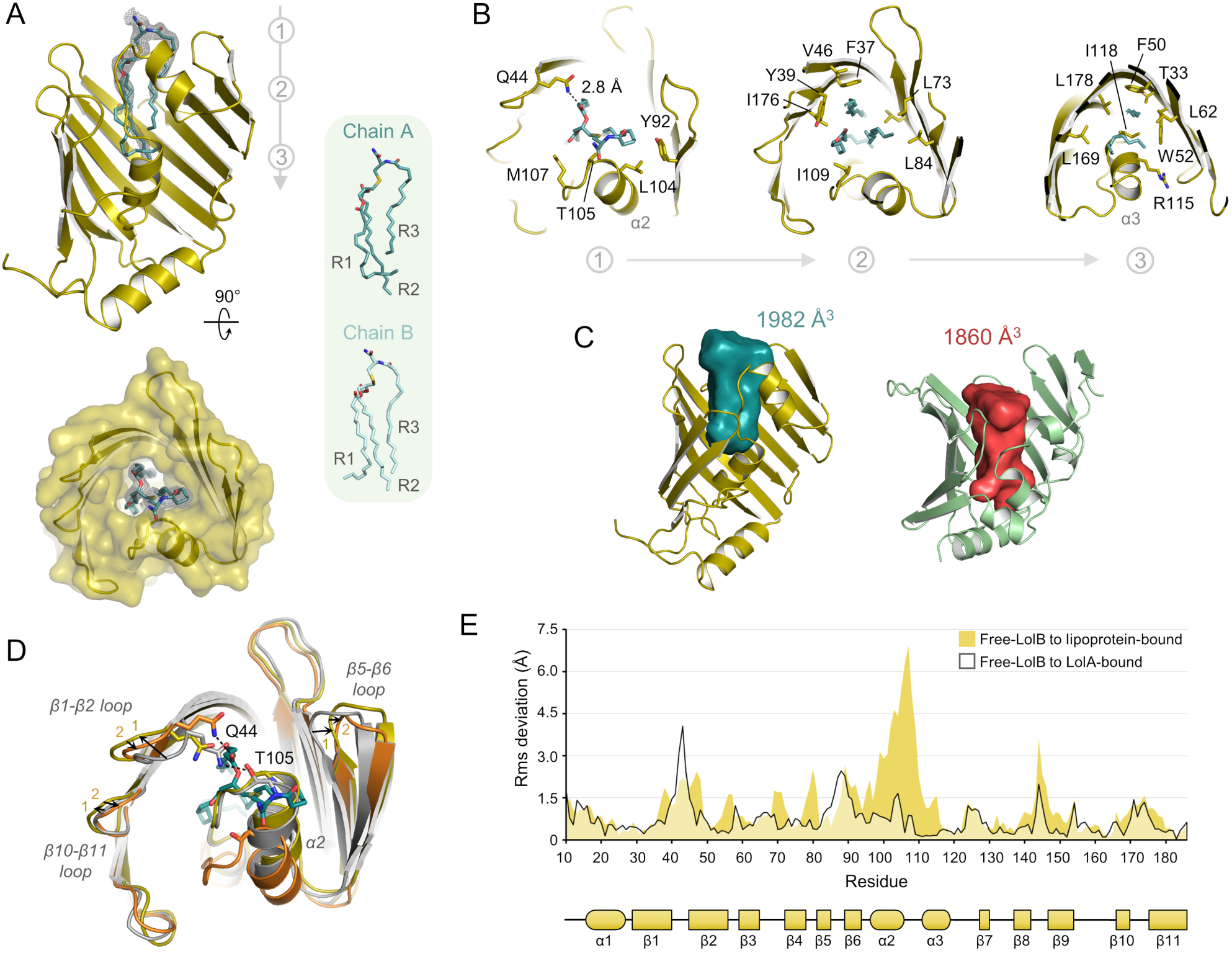
Structure of lipoprotein-bound LolB and conformation rearrangements upon association with LolA and lipoprotein. (A) Overall structure of LolB L114G (yellow) complexed with lipoprotein (cyan, stick representation). The mesh represents the Polder omit map of the lipoprotein ligand contoured at 3 σ. Positions of the lipoprotein acyl chains adopted in the two LolB chains within the asymmetric unit are displayed in green inset. (B) Cross section views of the LolB cavity cut at the points indicated in (A), displaying residues which interact with the lipoprotein acyl chains. (C) The LolB cavity visualized as a solid cyan cast in the LolB-lipoprotein and in the LolA-lipoprotein structure (7Z6W) with approximate volume indicated. (D) Alignment of LolB structures in its free (1IWM, light grey), LolA-bound (33DL, yellow) and lipoprotein-associated (33DM, orange) form. Arrows are highlighting movement adopted by indicated regions from the free to LolA-bound state 1) and from the latter to the lipoprotein-associated confirmation 2). The lipoprotein is shown in cyan. (E) Rmsd plots of residues from LolB in the free-state (1IWM) to LolA-bound (33DL, light grey) or to lipoprotein-associated conformation (33DM, yellow). LolB protein secondary structural elements are shown underneath.

**Figure 6.**
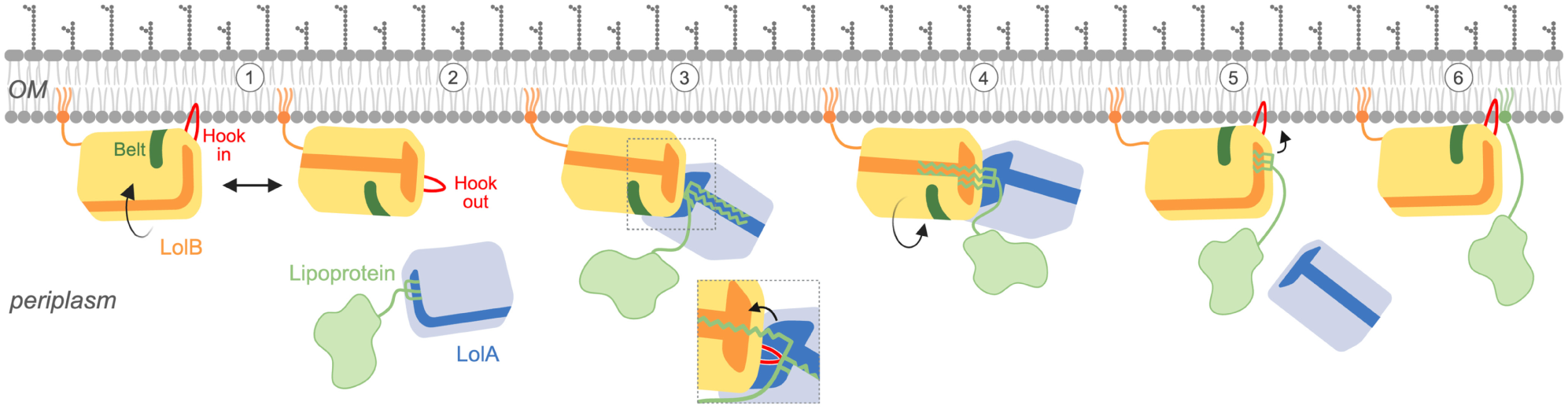
Model of lipoprotein transfer from LolA to LolB and subsequent membrane insertion. 1) Lipoprotein receptor LolB (yellow) can associate with the membrane in either a ‘Hook-in’ or ‘Hook-out’ conformation. The LolB Belt and Hook are highlighted in green and red, respectively. 2) LolA-lipoprotein complex approaches LolB. 3) In the ‘Hook-out’ conformation, LolB engages the LolA-lipoprotein complex via Belt residues, releasing the ‘latch’ of the LolB cavity entrance. Insertion of the LolB Hook into the LolA cavity initiates transfer of the lipoprotein acyl chains (inset). 4) Following transfer of the acyl chains into LolB, the protein rotates to a ‘Hook-in’ conformation. 5) LolA is released, and LolB initiates the insertion of the lipoprotein acyl chains into the membrane. 6) The lipoprotein is fully integrated into the outer membrane, leaving LolB free to engage a new LolA-lipoprotein complex.

We created a molecular morph to visualize the possible conformational changes undergone by LolB as it transitions from its unbound form to the LolA-associated and lipoprotein-complexed states (Movie S3). Association with LolA initially relaxes the mouth of the LolB cavity, with the loops joining β-strands 1-2, 5-6 and 10-11 shifting away from the α2 helix as shown in Figure 5D. A per residue rmsd plot indicates that these regions exhibit the largest displacements upon association with LolA (Figure 5E). This motion is likely induced by residues of the Belt (R34, R49, R175, K177 and K179), located on adjacent strands 1, 2 and 11, which move closer and create contacts with partner residues in LolA (Figure 2). This disrupts the Q44-T105 ‘latch’ hydrogen bond (Figure 5D and *SI Appendix* Figure 6B). As a result, the α2 helix shifts slightly toward the center of the cavity, breaking a secondary hydrogen bond between E98 (located at the base of the α2 helix) and Q143, positioned on the opposite side of the LolB cavity mouth (Movie S3 and *SI Appendix* Figure 6B). These conformational changes open the entrance of the cavity, priming LolB for lipoprotein transfer. We next compared the LolA- and lipoprotein-associated conformations of LolB (Figure 5E and *SI Appendix* Figure 6C). This analysis reveals that the most substantial motion occurs in the α2 helix and α2-α3 linker (residues 98-110) which shift by 2 to 7 Å away from the LolB β-barrel to fully open the LolB cavity. In contrast to LolA, where both α2 and α3 helices open to accommodate the lipoprotein (32), only the α2 helix of LolB shifts position while the α3 helix (residues 111 to 119) remains relatively stable. Arginine 115 consistently interacts with residues from the β2-β3 loop, anchoring the α3 helix to the rest of the structure across the three conformational states of LolB (Movie S3 and *SI Appendix* Figure 6B). LolB helices α2 and α3 are positioned closer to the cavity mouth than their counterparts in lipoprotein-loaded LolA. As a result, the cavity expansion is restricted to the upper half of the protein and the lipoprotein is inserted ∼ 11 Å less deeply inside LolB compared to LolA (*SI Appendix* Figure 4C). Structural alignment of the LolA and LolB lipoprotein-bound forms with our LolA-LolB complex reveals that, in both cases, the +1 cysteine is pointing away from LolA and LolB providing ample space for the flexible linker that connects the triacyl group to the lipoprotein globular domain. In this arrangement, the triacylated cysteine must translocate approximately 20 Å to reach LolB (*SI Appendix* Figure 6D), though we cannot exclude further rearrangements of LolA and LolB within the ternary complex that might reduce this distance to facilitate transfer.

### Alternate conformations of LolB on the membrane suggest a model for lipoprotein transfer from LolA and membrane insertion

Like other lipoproteins, LolB is tethered to the outer membrane via its triacyl group. Molecular dynamics simulations, however, suggest that the globular domain of LolB can directly associate with the membrane in two distinct conformations (45). In the primary binding mode, hereafter termed ‘Hook-in’, LolB lies against the membrane with all Belt residues, except R61, making contact with negatively-charged phospholipid headgroups (*SI Appendix* Figure 7A). In the secondary binding mode, here termed ‘Hook-out’, LolB is rotated by ∼90° around the cavity axis, exposing the protein Hook and Belt to the periplasm.

To provide further insight into how LolB receives lipoproteins from LolA and inserts them into the membrane, we aligned our LolA- and lipoprotein-bound structures of LolB with the ‘Hook-in’ and ‘Hook-out’ models. Structural alignment reveals that LolA can only interact with the ‘Hook-out’ conformation of LolB, in which the protein Hook and Belt are accessible, whereas the protein-protein interface is occluded by the membrane in the ‘Hook-in’ state (*SI Appendix* Figure 7B). In contrast, alignment with our lipoprotein-bound LolB structure suggests that the ‘Hook-in’ conformation is more favorable for lipoprotein release, as the acyl chains are positioned closer to the membrane surface.

We therefore propose a mechanism for how LolB engages LolA to receive lipoprotein and releases it into the outer membrane (Figure 6). Free LolB rests on the membrane lipid bilayer, alternating between a ‘Hook-in’ and ‘Hook-out’ conformation. LolA, bound to a lipoprotein, engages with the ‘Hook-out’ state of LolB, via interaction with Belt residues. This relaxes the entrance of the LolB cavity, breaking the Q44-T105 latch that maintains the helical lid closed, leaving LolB primed for acyl chain insertion. Concomitant insertion of the LolB Hook into the LolA cavity displaces the acyl chains, disrupting the bonds between LolA S150-Y152 and the lipoprotein R2 glycerol carbonyl (32). This initiates the transfer of the first acyl chain, most likely R3, toward LolB. The LolA-LolB structure suggests the lipoprotein acyl chains must translocate more than 20 Å. However, the LolB Hook may provide a conduit to facilitate the acyl chain transit and it is also possible that a reorientation of the two proteins brings both cavities closer. Once the acyl chains have been transferred, LolB rotates to the more stable ‘Hook-in’ conformation where the Belt residues interact with the negatively charged phospholipid headgroups, releasing LolA. The LolB cavity entrance is now located closer to the membrane, with the α2 helix pointing toward the periplasm. In this conformation, the Q44 residue is proximal to the phospholipid headgroups which could destabilize the interaction with the R2 acyl chain, allowing transfer into the membrane. The Hook, inserted into the membrane, forms a conduit for acyl chain transit, and potentially disrupts local phospholipid packing to aid insertion.

## Discussion

The final stages of lipoprotein trafficking depend on LolB, the outer membrane receptor that receives lipoproteins from the chaperone, LolA, and transfers them to the inner leaflet of the outer membrane. Here, we present a structure of LolB in complex with LolA (Figure 2) revealing that interaction between the two proteins is dependent on the LolB ‘Hook’, a β-hairpin loop reminiscent of that found in LolC (27), and ‘Belt’, a succession of charged residues spanning the LolB β-barrel. The two protein cavities adopt a conformation consistent with the previously proposed mouth-to-mouth model inferred from *in vivo* crosslinking data (*SI Appendix* Figure 1A), (40). However, instead of adopting a linear arrangement, the two proteins are oriented with an approximately 105° angle between their cavity axes, providing space for the lipoprotein linker to extend from the LolA-LolB complex.

A structure of *X. campestris* LolA-LolB (42) displays a similar conformation that is also dependent on a constellation of positively charged residues on the convex face of LolB. These residues are not conserved in primary sequence but create a similar network of interactions (*SI Appendix* Figure 1C) that is likely to be common in divergent LolA-LolB complexes.

Superimposition of our LolA-LolB complex with the lipoprotein-bound LolA structure (32) shows that the Hook apex residue, L68, clashes with one of the lipoprotein acyl chains inside LolA (Figure 3E). This suggests that insertion of the Hook initiates the lipoprotein transfer from LolA to LolB by physically disengaging the acyl chains from the LolA cavity. Consistent with this idea, altering the length of the Hook or modifying the nature of the apex residue, as performed here (Figure 3) and by Hayashi and colleagues (36), dramatically affects the ability of LolB to accept lipoprotein.

The hydrophobic cavities of LolA and LolB are occluded by a helical lid which moves out to accommodate the lipoprotein cargo. In LolA, this lid is secured at the base of the cavity by R43 (41) but LolB lacks an equivalent residue. However, by targeting residues that mediate interactions between the helical lid and the core structure of the protein, we identified residue Q44 which is located at the entrance of the cavity and forms hydrogen bonds with two residues from the LolB helical lid. Mutation of Q44 to alanine resulted in fivefold greater cavity volume (Figure 4) demonstrating that Q44 acts as a molecular latch maintaining the LolB cavity in a closed state.

We previously demonstrated that the LolC Hook opens the LolA cavity, priming the chaperone to receive lipoprotein (27). Similarly, the interaction of LolA with LolB induces conformational changes sufficient to break the Q44 latch (*SI Appendix* Figure 6B), leaving LolB poised for lipoprotein insertion. To date, structures of both free and LolA-complexed LolB are available only for *E. coli* proteins. Further work is therefore required to determine whether this mechanism is conserved in bacteria containing both LolA and LolB homologues.

Finally, we solved the structure of lipoprotein complexed LolB L114G (Figure 5), a variant of LolB that receives lipoprotein more efficiently than wild-type LolB but fully supports *in vivo* function. Like LolA, LolB accommodates all three acyl chains within its cavity but notably, the acyl chains are less deeply inserted in LolB (*SI Appendix* Figure 6C). This observation aligns with the distinct roles of the two proteins; LolB accepts and presumably inserts lipoproteins into the membrane rapidly, obviating the need for a stable, long-lived complex and potentially facilitating lipoprotein diffusion into the membrane. In contrast, the LolA-lipoprotein complex must traverse the periplasm to reach LolB, necessitating a greater stability. In both LolA and LolB, interactions of the hydrophobic residues lining the cavity with the acyl chains are a major contributor to lipoprotein binding. However, specific residues make hydrophilic interactions with the lipoprotein carbonyl groups. In LolA, residues S150 and Y152 form hydrogen bonds with the carbonyl of chain R2 (32) and both also interact with the backbone carbonyl of LolB L68, demonstrating that Hook insertion likely disrupts these key stabilizing bonds to facilitate lipoprotein transfer. Similarly, LolB Q44 which helps maintain the lipoprotein lid in a closed state also directly interacts with the R2 carbonyl group in the lipoprotein-LolB structure, possibly guiding the insertion of the lipoprotein into the LolB cavity during transfer from LolA.

LolB is attached to the outer membrane via its N-terminal triacyl group but this anchor is not essential for function (33). This is consistent with molecular dynamics simulations (45) showing that the globular domain of LolB can interact with the membrane in two conformations: a ‘Hook-in’ and a ‘Hook-out’ state, based on the Hook proximity to the membrane. Structural superposition of our LolA- and lipoprotein-bound LolB complexes reveals that only the Hook-out conformation allows LolA association, whereas the Hook-in state is more favourable for lipoprotein release into the membrane (Figure 6). The LolB Hook is therefore important not only for lipoprotein transfer from LolA but also for membrane insertion, as previously suggested (36).

The structural, biochemical and *in vivo* data presented here provide more insight into the final steps of the Lol pathway for bacteria which encode discrete LolA and LolB proteins. The insertion mechanism proposed here is also likely to apply to those bacteria expressing a single LolA/B homologue since an extended Hook-like loop is important for function of these bifunctional proteins (37). Our model for the LolA-LolB system provides a rationale for release of the LolA chaperone but further experiments are required to understand how the bifunctional LolA is released from the membrane following lipoprotein transfer.

The structures resolved here provide a basis for inhibiting the Lol pathway by targeting LolB complexes with small molecules and would augment efforts to interfere with the action of LolCDE (23, 24, 26, 46). Virtual screening identified the first small molecules that inhibit *Vibrio parahaemolyticus* growth and bind to LolB *in vitro* (25). Computational docking suggested the molecule interacts with several residues including a threonine located on the helical lid and a glutamine residue at the entrance of the cavity. By analogy to our structure these residues would form a molecular latch in *Vibrio parahaemolyticus* LolB and interaction with the small molecule could inhibit LolB by destabilizing the closed state. Our structures provide a basis for the identification of other LolB specific inhibitors that could for example block the LolA-LolB association or lipoprotein transfer and could complete efforts made to inhibit the Lol pathway. As LolB is biosynthetically more expensive to produce and is itself a substrate of the Lol system, molecules with conceptually similar activities may be more effective than targeting LolA.

In summary, our results provide insight into the molecular basis for chaperone and lipoprotein recognition by the outer membrane receptor LolB and provide a key step towards the complete understanding of a fundamental trafficking pathway.

## Methods

### Construction of strains and plasmids

Primer sequences and strains/constructs used in this study are detailed in Tables S2 and S3, respectively. To construct an arabinose-inducible *lolB* conditional knockout strain (BW65), the *lolB* locus including the ribosome binding site was amplified using primers P1 and P2 and cloned into the NheI-HindIII sites of pBAD18 (47). The region encompassing the *araC* gene, pBAD promoter, *lolB* and the downstream terminator was amplified using primers P3 and P4, digested ClaI-SacI and cloned into the integration vector pLDR9 digested with the same enzymes. The construct was integrated into the lambda *attB* site of *E. coli* BW25113 according to a previously described protocol (48). The native copy of *lolB* was replaced with a kanamycin resistance cassette by amplifying pKD13 with primers P5 and P6 using the λ Red recombinase system as described (49) except that pSIM5 (50) was used for recombinase expression. Deletions were confirmed by PCR of the gene locus with primers P7 and P8.

To enable complementation of BW65, plasmid pCDF-LolB_alfa_ was constructed. A fragment encoding full-length *lolB* with a C-terminal Alfa tag (51) was synthesized (IDT DNA), amplified with primers P9 and P10, and introduced by Gibson assembly (52) into pCDFDuet digested with NcoI and KpnI. Plasmids pET28-LolB and pET28-LolA, were used to produce N-terminally His-tagged non-acylated LolB and mature LolA proteins respectively and have been described previously (27). Mutations were introduced into pCDF-LolB_alfa_ and pET28-LolB by Quikchange site-directed mutagenesis (Agilent) using primers listed in Table S2. To enable TEV mediated removal of the His-tag on LolB, a gene fragment encoding an N-terminal octahistidine tag, TEV cleavage site and mature LolB (residues 23-207) was synthesized (IDT DNA), digested with NcoI and BamHI and ligated into pET28 digested with the same enzymes resulting in pET28-LolB_TEV/His_. Subsequently, a L114G mutation was introduced by Quikchange (Agilent). Creation of plasmids pCDF-LolA-Pal_WT(OCTA)_ and pCDF-LolA_Strep_-Pal_TEV/FL2_ has been previously described (32). All clones were verified by DNA sequencing (Source Bioscience).

### Complementation of LolB variants

*E. coli* strain BW65 bearing pCDF LolB or the indicated variant was cultured in LB medium supplemented with 0.2% arabinose and appropriate antibiotics. The next day, 1 mL of overnight culture was centrifuged at 7,000 *g* for 2 mins, washed twice in LB supplemented with 0.1% D-fucose and then diluted 1/100 into fresh LB containing 0.1% fucose. Cells were then grown to an OD_600_ of 0.5, collected at 7,000 *g*, washed twice in LB and then serial tenfold dilutions performed in LB. Dilutions were plated out on LB agar with and without arabinose and grown overnight at 37 °C before imaging the next day. To assess expression level of the variant proteins, whole cell samples were resolved on 12% SDS-PAGE gels, transferred to PVDF membrane, and immunoblotted with a 680 nm IR dye conjugated anti-Alfa antibody (Nanotag). Immunoblots were revealed using an Odyssey Licor fluorescence imager.

### Protein purification

#### Purification of LolA and LolB

*E. coli* BL21 (DE3) cells bearing plasmid pET24-LolA, pET28-LolB_His_, pET28-LolB(L114G)_TEV/His_, or appropriate variant were grown in 1L 2YT medium supplemented with 50 µg/µL kanamycin at 37 °C and agitated at 200 rpm. This allows over-expression of a soluble, non-acylated LolB in the cytoplasm. When the culture reached an OD_600_ of 0.6-0.8, the temperature was reduced to 18 °C and protein expression was induced with 0.1 mM isopropyl β-D-1-thiogalactopyranoside (IPTG) for 18 hours of further growth. Cells were harvested by centrifugation at 6,000 *g* for 6 min and the pellet was resuspended in buffer 50 mM Tris pH 7.5, 300 mM NaCl with EDTA-free protease inhibitors (Roche), 0.1 mg/mL lysozyme (Sigma) and 10 µg/mL DNase (Sigma). Bacteria were lysed by cell disruption at 32,000 psi (Constant Systems) before removal of cell debris by ultra-centrifugation for 1.5 hours, 190,000 *g* at 4°C. A final concentration of 20 mM imidazole was added to the clarified lysate before loading onto a 5 mL HisTrap FF column (GE Healthcare) using an AKTA Xpress system. The column was washed with 15 column volumes (CV) of the same buffer containing 20 mM imidazole, before elution with 250 mM imidazole. Proteins were dialyzed against 2 L of 25 mM HEPES pH 7.5, 150 or 200 mM NaCl for LolA and LolB respectively, using 8,000 Da MWCO dialysis tubing (Fisher Scientific) overnight at 4°C. After concentration to ∼30 mg/mL using 10,000 Da MWCO protein concentrator (Amicon), proteins were flash frozen in liquid nitrogen and stored at -80°C. Where necessary, His-tags from LolA, LolB and variants were removed using a thrombin CleanCleave Kit (Sigma) following the manufacturer’s instructions.

#### Purification of LolA-Pal complexes

*E. coli* C43 cells bearing plasmid pCDF-LolA-Pal_WT(octa)_, pCDF-LolA-Pal_TEV/FL2_ or pCDF-LolA(R43L)_His_-Pal were grown in 2YT medium supplemented with 0.5% glycerol and 50 µg/µL streptomycin at 37°C with agitation at 200 rpm. When the culture reached an OD_600_ of 0.6-0.8, expression was induced with 1 mM IPTG. After a further 2 hours growth, cells were harvested by centrifugation at 3,250 *g* for 20 minutes. Spheroplasts were produced by resuspending the bacteria in a buffer composed of 200 mM Tris pH 8.0, 500 mM sucrose, 1 mM EDTA and 0.5 mg/mL lysozyme. After incubation at room temperature for 60 minutes, cells were centrifuged at 20,000 *g* for 30 minutes to remove spheroplasts and intact cells. The periplasmic fraction obtained was subsequently centrifuged at 190,000 *g* for 45 minutes. The supernatant was then supplemented with 5 mM MgCl_2_, 300 mM NaCl and 20 mM imidazole before loading onto a 5 mL HisTrap FF column. The resin was washed and bound proteins eluted with the same buffer containing 250 mM imidazole. When required, His-tags were removed by incubation with 30 μg TEV protease (53) per mg of LolA-Pal complex overnight at 4°C before IMAC mediated removal of His-tags, uncleaved proteins and His-tagged TEV protease. Proteins were dialyzed into 25 mM HEPES pH 7.5, 200 mM NaCl, concentrated to ∼10 mg/mL and flash frozen in liquid nitrogen.

### Production of protein complexes

#### LolA R43L-LolB

LolA R43L_his_-Pal was incubated with tag-free, non-acylated LolB in a molar ratio of 1:3 for 30 minutes in 25 mM HEPES pH 7.5, 150 mM NaCl, before incubation for 15 minutes with 1 mL Profinity nickel resin (50%, BioRad). The resin was then washed with 30 mL buffer, proteins eluted with 250 mM imidazole and dialyzed against 2 L of 25 mM HEPES pH 7.5, 200 mM NaCl using 8,000 MWCO dialysis tubing (Fisher Scientific) overnight at 4°C. Proteins were concentrated, snap frozen in liquid nitrogen, and stored at -80 °C.

#### LolB lipoprotein complexes

LolA-Pal complexes produced from pCDF-LolA_strep_-Pal_TEV/FL2_ contain a Pal with a TEV cleavage site was introduced at position 49 of full-length Pal (position 28 in the mature sequence) facilitating removal of the mature domain of Pal (32). These complexes were mixed with LolB-L114G_TEV/His_ in a molar ratio of 5:1 to maximize transfer of Pal. The mixture was incubated at 37°C for up to 5 hours. The protein mix was incubated with Profinity nickel resin (BioRad) which was washed extensively with 25 mM HEPES pH 7.5, 150 mM NaCl to remove free LolA_strep_ before eluting His-tagged proteins with 250 mM imidazole. The elution fractions were concentrated and then buffer exchanged into 100 mM Tris pH 8.0, 150 mM NaCl, 1 mM EDTA, before incubation with Strep-Tactin XT resin (IBA) to bind remaining LolA_strep_ and LolA_strep_-Pal_TEV/FL2_. The resin flow-through and wash fractions were pooled, concentrated and 30 µg TEV protease per mg of protein added to remove the mature domain of Pal, overnight at 4°C in the presence of 2 mM TCEP. The next day, 5 mM MgCl_2_ was added to the mixture before removing His-tagged cleaved proteins and TEV proteases by IMAC. The resin flow-through and wash fractions which contain LolB-lipoprotein were buffer exchanged into 25 mM HEPES pH 7.5, 150 mM NaCl, concentrated, snap frozen in liquid nitrogen, and stored at -80 °C.

### Protein crystallization and structure determination

All crystallography trials were set up using the liquid handling robot Mosquito (TTP) and performed at 15 °C using the sitting-drop vapor-diffusion method over a reservoir of 80 µL in MRC 2-drop plates (Molecular Dimensions). Prior to being frozen in liquid nitrogen, crystals were cryoprotected in the reservoir solution supplemented with 20% glycerol except for the LolA-LolB complex crystal which grew in a solution already containing 40% glycerol ethoxylate. Diffraction and refinement statistics can be found in Table S1.

#### LolA R43L-LolB

The LolA R43L-LolB complex at 7.7 mg/mL was mixed at a protein:reservoir ratio of 1:1 in 1 µL final volume. Crystals were obtained in 100 mM sodium HEPES pH 7.5, 20% PEG 4000 and 10% 2-Propanol (Structure Screen, Molecular Dimensions) and diffracted remotely on beamline I04 at Diamond Synchrotron. Diffraction data were processed using the CCP4 program suite (54). Images were first indexed and integrated with iMosflm (55). Data were then scaled and merged with Aimless (56) and data truncated at an appropriate resolution. Molecular replacement was performed with Phaser (57) using wild-type LolB (PDB 1IWM) and LolA (PDB 1UA8) as search models. The asymmetric unit contains one molecule of LolA and one of LolB The structure was refined with cycles of manual building with Coot (58), and restrained refinement with Phenix (59).

#### LolB L114G-lipoprotein

Crystals of LolB L114G complexed with lipoprotein were obtained in 40% glycerol ethoxylate (Midas plus, Molecular Dimensions) at a protein:reservoir ratio of 1:1 in 1 µL final volume. The structure was solved as described for the LolA-LolB complex by molecular replacement with Phaser using LolB (PDB 1IWM) as a search model. and data collected at Diamond I24. Density inside the cavity of LolA enabled the modeling of the three lipoprotein acyl chains and corresponding triacylated cysteine but not for the remaining residues. The acyl chains were modeled as C16 consistent with the majority acyl chain length in *E. coli* lipoproteins (60). Two slightly different conformations of the terminal six carbon atoms of acyl chain R1 chain were evident in each of the two copies of the complex present in the asymmetric unit. The structure was refined with cycles of manual building with Coot (58), and restrained refinement with Refmac (61).

#### LolB Q44A

Purified non-acylated protein was concentrated to 15 mg/mL and crystal trays set up using an ‘in-house’ designed screen based on the crystallization condition described previously for wild-type LolB (34). The closed LolB Q44A structure was obtained from a crystal formed in 30% PEG 2000 MME, 100 mM sodium acetate pH 5.5, 200 mM ammonium sulfate, and 0.15 mM sodium iodide. Seeding from an identical condition was performed. The LolB Q44A open form was solved from a crystal which crystallized in 39% PEG 2000 MME, 100 mM sodium acetate pH 6.2, and 200 mM ammonium sulfate. In that case, the protein solution also contained 1 mM 11-(dansylamino) undecanoic acid (DAUDA, Cayman Chemical), however no density for the molecule was observed. Diffraction data were collected at Diamond I04 or I04-1, integrated and scaled with XDS (62). Structures were solved as described for the LolB-lipoprotein complex using LolB (PDB 1IWM) for molecular replacement and contain one molecule per asymmetric unit. TLS groups from Refmac were used during refinement.

All the final structures were validated with Rampage (63) and Procheck (64)

### Analysis of protein cavities

The protein cavities for wild-type LolB (1IWM), the open and closed LolB Q44A, the lipoprotein-bound LolB L114G and the LolA-lipoprotein complex (7Z6W) structures were visualized using HOLLOW (65) and the cavity volumes determined with 3V (66).

### Interaction between LolA and LolB

His-tagged soluble wild-type or variant LolB (15 µM) was incubated with 15 µM untagged wild-type LolA wild-type, in a final volume of 250 µL buffer (25 mM HEPES pH 7.5, 150 mM NaCl). Samples were incubated at room temperature (∼22°C) for 5 minutes before loading on 100 µL pre-washed Ni-IMAC slurry (BioRad) and incubating for a further 5 minutes. Mini spin columns (Generon) were used to separate resin from supernatant. The resin was washed three times with 500 µL buffer before elution of bound proteins with buffer containing 250 mM imidazole. Eluted proteins were analyzed on SDS-PAGE gels before scanning and quantification with an Odyssey CLx and ImageStudio software (Licor). Protein band intensity was measured and recorded as LolA per LolB, and performance of the LolB mutants was normalized against the mean results for wild-type LolB. Assays were performed in triplicate and mean results were presented with standard deviation.

### Transfer of lipoproteins from LolA to LolB

LolA-Pal complexes from which the His-tag had been cleaved were mixed at a 1:1 ratio with 15 µM His-tagged soluble LolB in a final volume of 250 µL buffer containing 25 mM HEPES pH 7.5, 200 mM NaCl. After 5 minutes on ice, the mixtures were incubated with 100 µL of Ni-IMAC resin (Biorad) for 5 minutes in microbatch spin columns (Generon). The resin was washed three times with 500 µL of ice-cold buffer prior to elution of bound proteins with 250 µL buffer containing 250 mM imidazole. The elution fractions were analyzed on 12% SDS-PAGE gels including purified LolB and LolA-Pal complexes as references.

### Fluorescence spectroscopy

Fluorescence measurements of the fluorescent fatty acid probe 11-(dansylamino) undecanoic acid (DAUDA, Cayman Chemical) were performed in a FluoroLog (Horiba) spectrometer at room temperature in a final volume of 1.2 mL buffer (20 mM HEPES pH 7.5, 150 mM NaCl) using a 2 ml quartz cuvette. DAUDA stock solution was prepared at a concentration of 10 mM in DMSO and used at a final concentration of 10 µM. Purified soluble wild-type or variant LolB protein (40 µM final concentration) or an equivalent volume of buffer was then added, and fluorescence emission was recorded between 400 and 600 nm after excitation at 335 nm.

## Acknowledgements

We acknowledge Diamond Light Source for time on beamlines I04, I04-1 and I24 and support from the CNRS/IN2P3 Computing Center (Lyon - France) for providing computing and data-processing resources. We also thank Susanne van den Berg for the gift of plasmid pTH24:TEVSH. This work was supported by grants from the UK Medical Research Council (MR/N000994/1 and MR/V000616/1) and a starting grant to E. K. from the CNRS National Institute of Biological Sciences (INSB 266416/266417). We are thankful to the Department of Pathology, University of Cambridge (UK), for granting V. K. a PhD studentship to support A. E. J.

## Data availability

Coordinates and structure factors were deposited in the Protein Data Bank under accession codes **33DL** (LolA R43L-LolB), **33DM** (LolB L114G-lipoprotein), **33DN** (LolB Q44A-open form) and **33DO** (LolB Q44A-closed form).

## Competing Interest Statement

The authors declare no conflict of interest.

## Supplementary Information

**Supplemental Figure 1.**
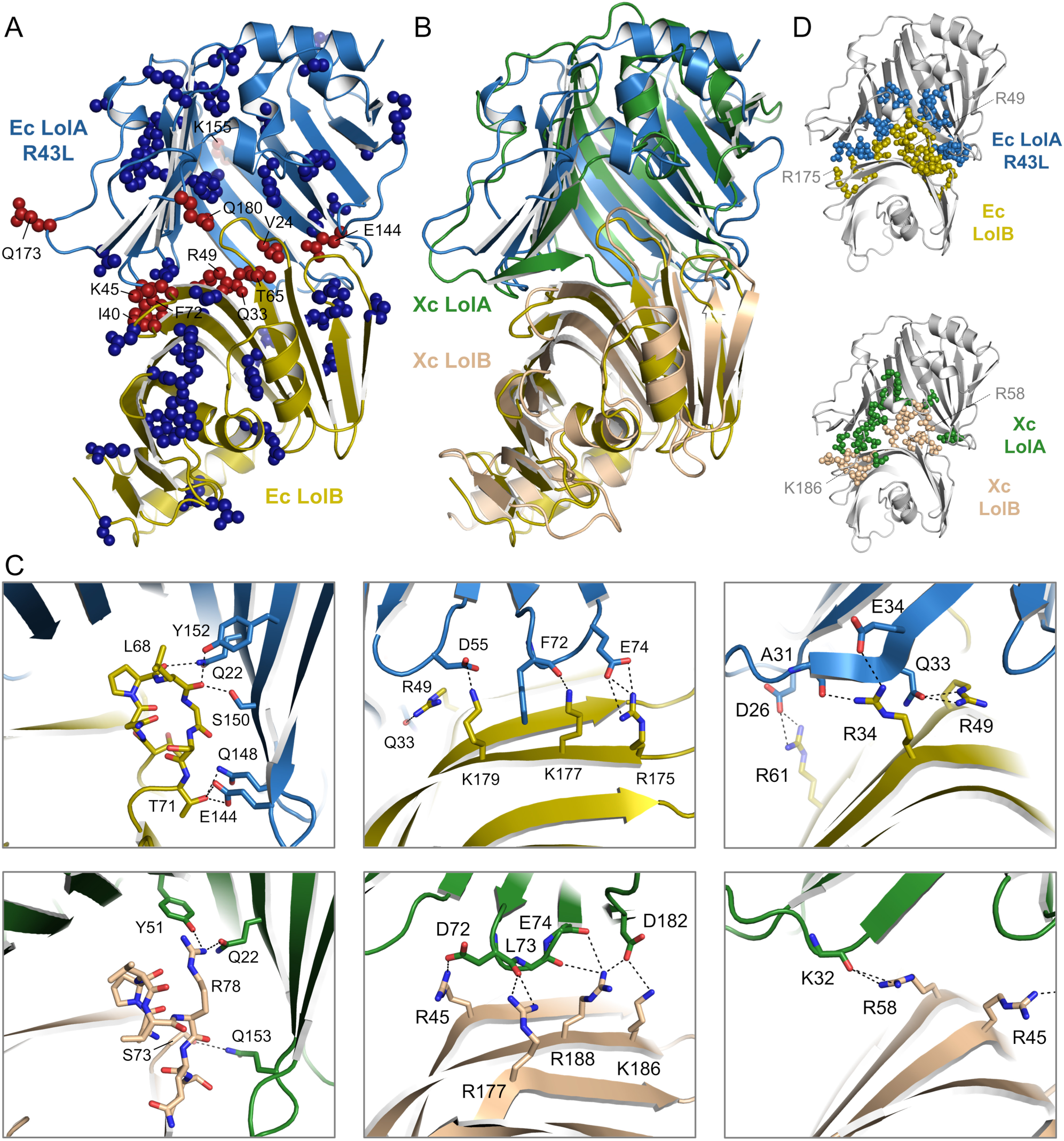
Comparison of *E. coli* LolA R43L-LolB structure with published crosslinking and structural data. (A) LolA and LolB *in vivo* photo-crosslinking results (40) are mapped onto our LolA R43L-LolB structure. Residues which formed photo-inducible crosslinks when replaced with pBPA (p-benzoyl-L-phenylalanine) are shown in red, whereas residues unable to crosslink are in blue. (B) Alignments of *E. coli* LolA R43L-LolB (Ec) with the published *X. campestris* (Xc) LolA-LolB structure (8ORN) (42). (C) Close-up views of LolB Hook and Belt residues in the *E. coli* and *X. campestris* structures. (D) Residues located at the protein-protein interface in the LolA-LolB structures are shown as spheres (*E. coli* 33DL and *X. campestris* 8ORN).

**Supplemental Figure 2.**
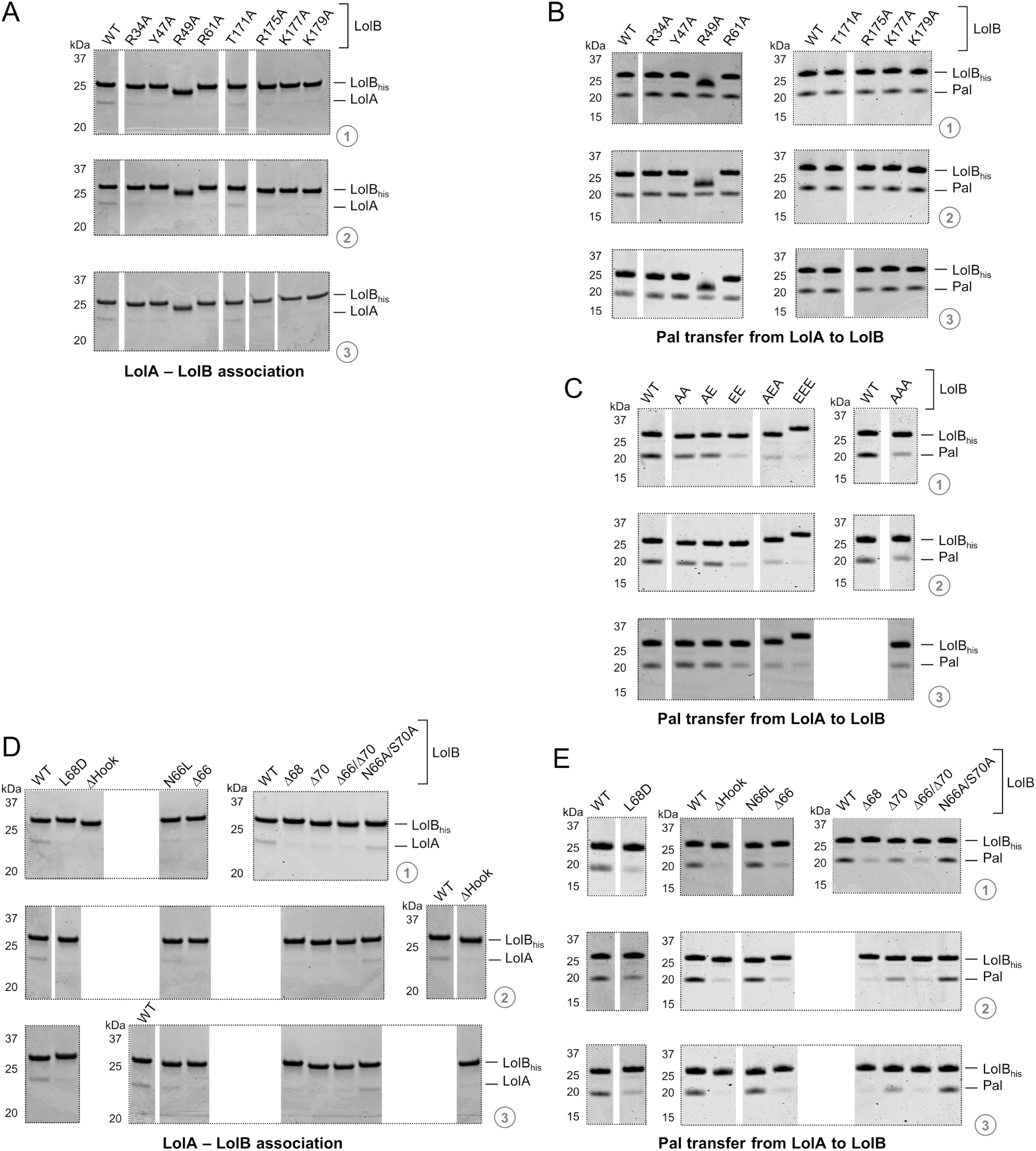
Raw data for the LolA-LolB association and lipoprotein transfer assays presented in. **Figure 3**. (A) SDS-PAGE images for triplicate repeats of the *in vitro* interaction assay of LolA and wild-type LolB (WT) or the indicated variants quantified in Figure 3A. His-tagged LolB proteins were incubated with tag-free LolA prior to immobilization on Ni resin. After washing, bound proteins were eluted and analyzed by SDS-PAGE. (B) SDS-PAGE images for triplicate repeats of the *in vitro* assay of lipoprotein transfer from LolA to wild-type LolB (WT) or the indicated variants quantified in Figure 3A. His-tagged LolB proteins were incubated with LolA-Pal complex prior to immobilization on Ni resin. After washing, bound proteins were eluted and analyzed by SDS-PAGE. (C) Triplicate gel images for the Pal lipoprotein transfer assay for variants quantified in Figure 3C; data were obtained as described in (B). (D) Images of the triplicate repeats of the LolA-LolB *in vitro* interaction assay for the variants quantified in Figure 3F; data were obtained as described in (A). (E) Images for the triplicate repeats for the Pal lipoprotein transfer assay for variants quantified in Figure 3F; data were obtained as described in (B).

**Supplemental Figure 3.**
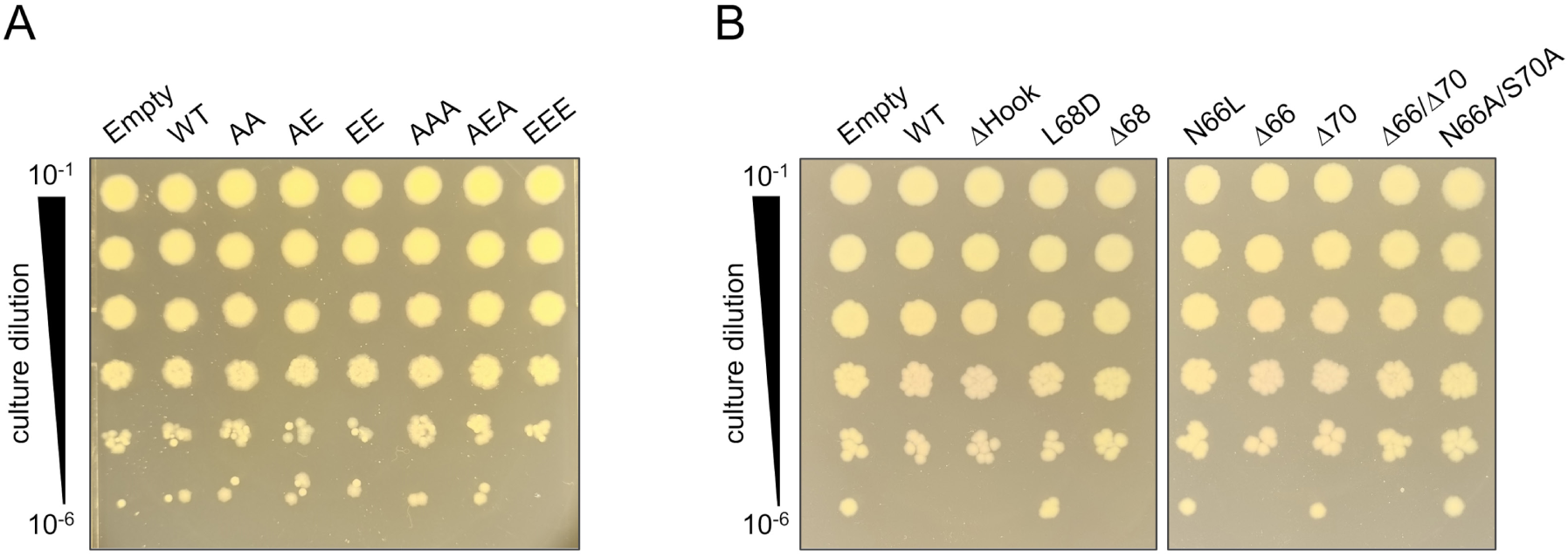
Control plates for the *in vivo* characterization of LolB Belt and Hook mutants. (A) Serial dilutions of a conditional *lolB* knockout *E. coli* strain (BW65) carrying plasmid-borne wild-type *lolB* or indicated variants in the R34/R61 (RR) or R175/K177/K179 (RKK) Belt cluster. Cells were grown in the presence of 0.2% arabinose to induce expression of chromosomally encoded wild-type *lolB*. The plate is a control for the experiment shown in Figure 3D. (B) Serial dilutions of *E. coli* BW65 carrying plasmid-borne wild-type *lolB* or indicated Hook variants in the presence of 0.2% arabinose to induce expression of chromosomally encoded wild-type *lolB*. The plate is a control for the experiment shown in Figure 3G.

**Supplemental Figure 4.**
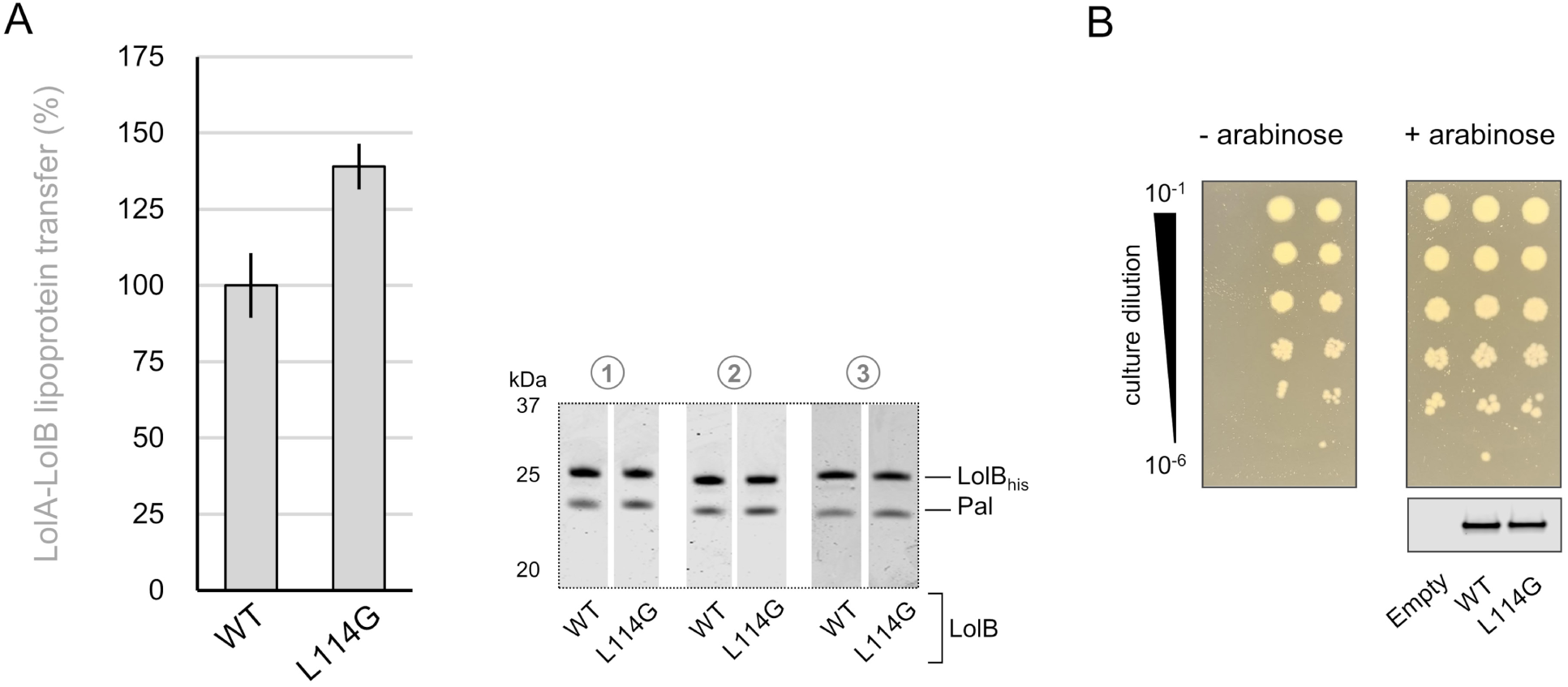
Characterization of the LolB L114G mutant. (A) His-tagged LolB wild-type or L114G variant were incubated with untagged LolA-Pal complexes. LolB and associated Pal were isolated on Ni resin and the elution fraction loaded onto SDS-PAGE. The amount of Pal transferred to LolB is reported, normalized to the value for wild-type LolB. Data shown are mean values ± standard deviation from triplicate experiments with raw gel images shown on the right for each repeat. (B) Serial dilutions of a conditional *lolB* knockout *E. coli* strain carrying either plasmid-borne wild-type *lolB* or L114G variant in the absence (left) or presence (right) of inducer required for expression of chromosomal *lolB* (top). Immunoblot showing the expression of plasmid-borne wild-type *lolB* and L114G variant in *E. coli* supported by growth of chromosomal *lolB* (bottom).

**Supplemental Figure 5.**
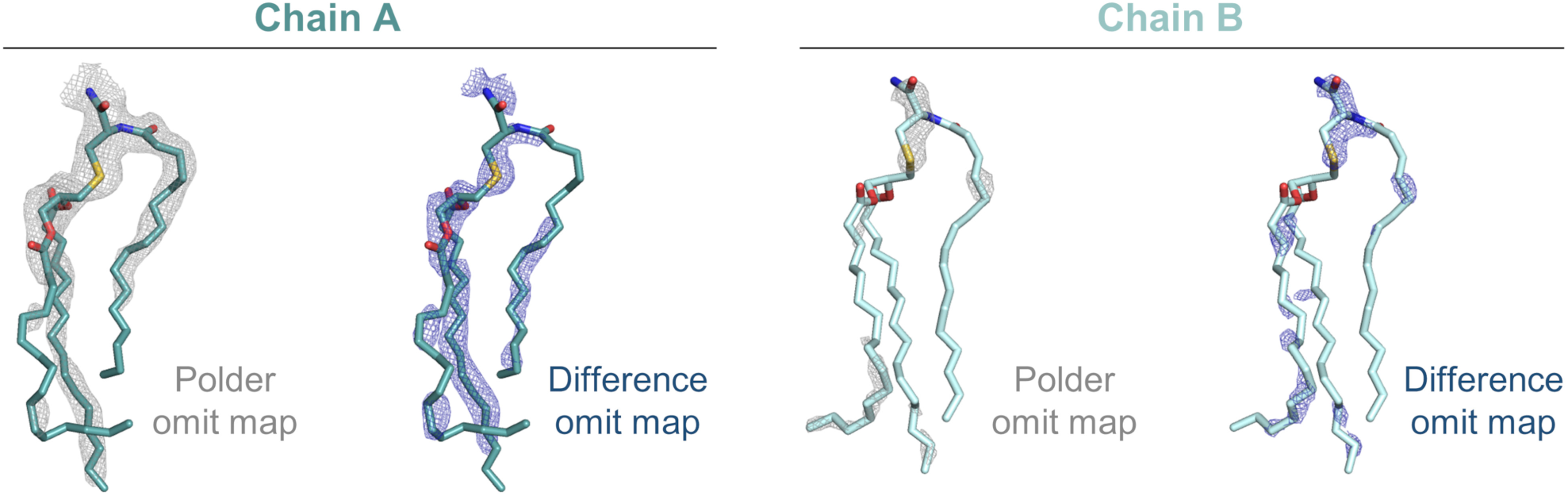
Polder and difference omit maps for the triacyl ligand in the LolB L114G lipoprotein structure. The triacyl group ligands in chain A (left, teal) and chain B (right, cyan) are shown in stick representation. For each chain, the Polder omit map and the *F_o_*-*F_c_* difference densities, both contoured at 3 σ, are represented as grey and blue meshes, respectively.

**Supplemental Figure 6.**
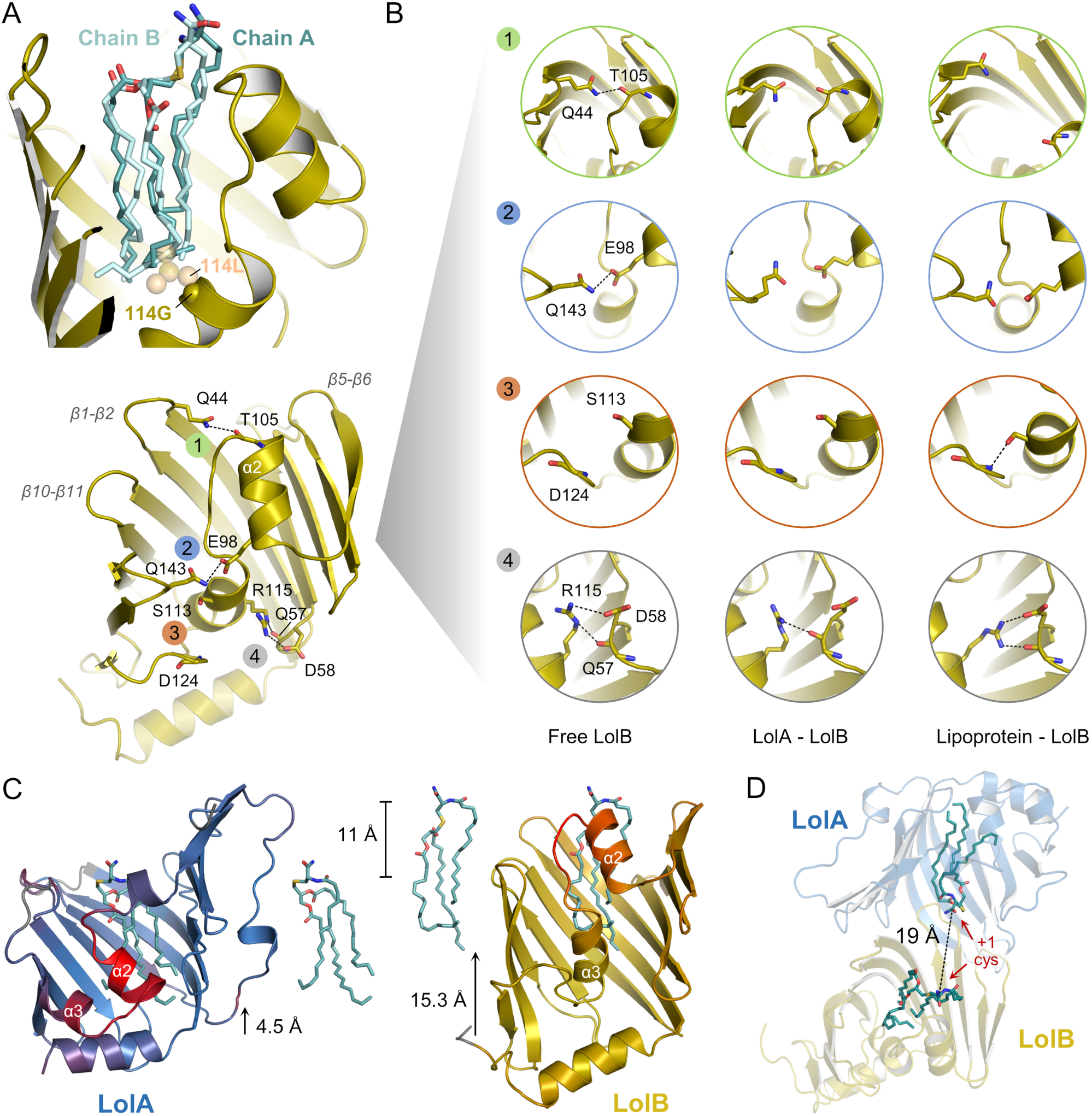
Structural comparison of LolB L114G in its free, LolA-bound and lipoprotein-associated form. (A) Superimposition of the two lipoprotein ligands present in the asymmetric unit of the lipoprotein-bound LolB L114G crystal. The position of G114 C_α_ is indicated by a yellow sphere. The native L114, modeled in the lipoprotein-bound structure, is shown as orange transparent spheres. (B) Major conformational changes of LolB from its free form to the LolA- and lipoprotein-associated state. Circles represent close-up views of the indicated regions of LolB (left, 1IWM) in each state. A molecular morph highlighting the LolB transition between these three states is shown in Movie S3. (C) The residue-by-residue rmsd between free and lipoprotein-associated LolA (1UA8, 7Z6W) or LolB (1IWM, 33DM) is displayed on the lipoprotein-bound LolA (7Z6W, blue) or LolB (33DM, yellow) structures. Rmsd values (Å) per aligned C_α_ atom are shown as a continuous gradient from blue or yellow to red with a maximal deviation of 8 Å. Distance between the C-terminus of α1 helix at the bottom of the cavity and the closest lipoprotein atom is indicated, together with the relative lipoprotein position in the LolA and LolB structures. (D) Alignment of lipoprotein-bound LolA (7Z6W, blue) and LolB (33DM, yellow) onto our LolA-LolB structure. The distance between the invariant +1 cysteine from each lipoprotein (cyan) is indicated.

**Supplemental Figure 7.**
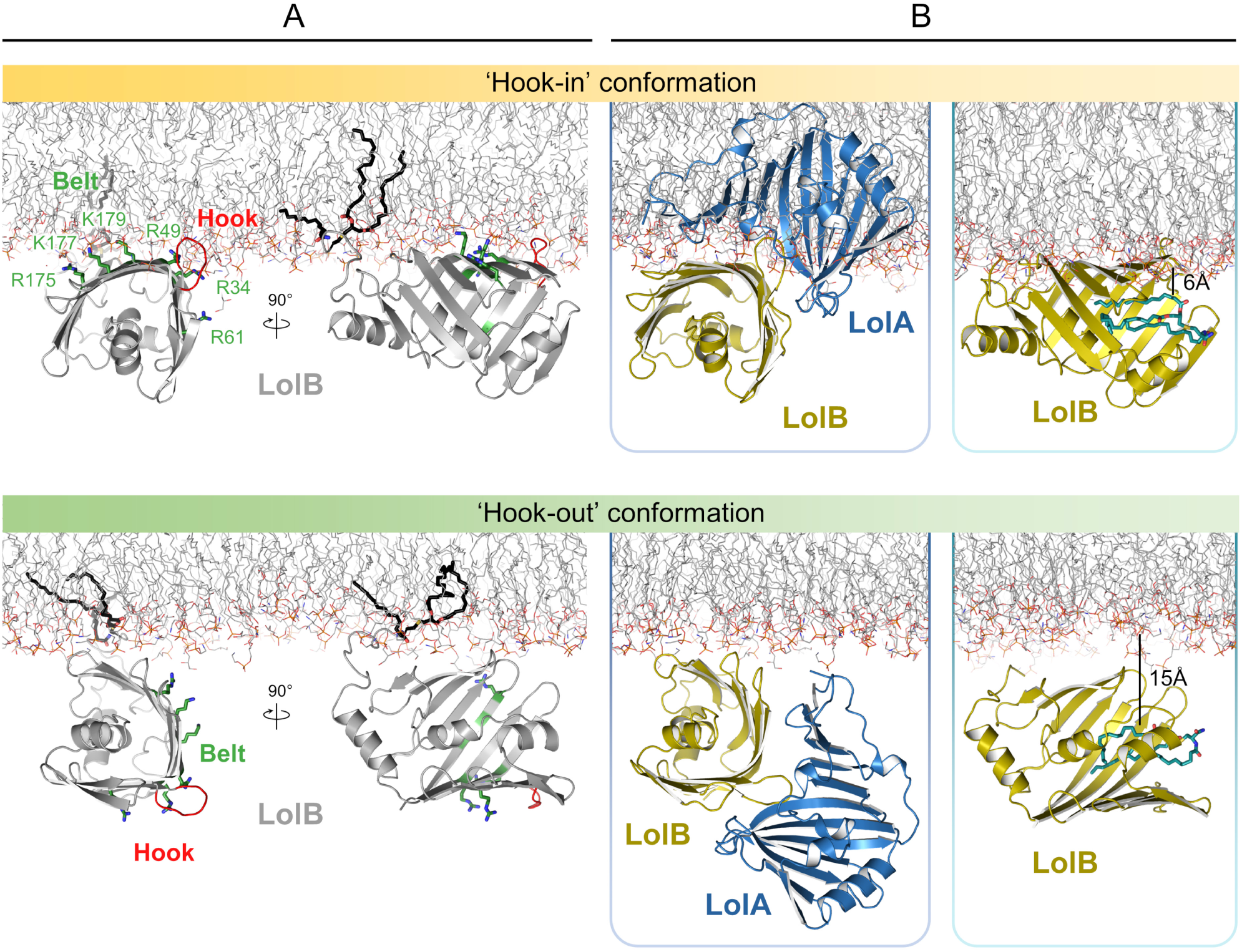
Comparison of simulated states of LolB on the membrane and their superimposition with LolA- and lipoprotein-bound structures. (A) Molecular dynamics derived ‘Hook-in’ and ‘Hook-out’ models of LolB associated with the membrane (45). LolB Hook is in red and residues from the Belt are represented as green sticks. The acyl chains of LolB are shown in black. (B) Superimposition of the LolA R43L-LolB (left) and lipoprotein-bound LolB L114G (right) structures with the ‘Hook-in’ and ‘Hook-out’ models of LolB derived from molecular dynamics simulations. The lipoprotein is shown in cyan with its approximate distance to the membrane indicated.

## Supplementary Tables

**Table S1.** X-ray data and refinement statistics.

|  | **LolB - LolA R43L complex** | **LolB L114G - lipo-protein complex** | **LolB Q44A (open form)** | **LolB Q44A (closed form)** |
|---|---|---|---|---|
|  | *(33DL)* | *(33DM)* | *(33DN)* | *(33DO)* |
| **Data Collection** |  |  |  |  |
| Beamline | Diamond I04 | Diamond I24 | Diamond I04-1 | Diamond I04 |
| Wavelength (Å) | 0.9795 | 0.9999 | 0.9159 | 0.9795 |
| **Crystal parameters** |  |  |  |  |
| Space Group | P2 2<sub>1</sub> 2<sub>1</sub> | P2<sub>1</sub> 2<sub>1</sub> 2<sub>1</sub> | P6<sub>3</sub> 2 2 | P6<sub>3</sub> 2 2 |
| Unit Cell Dimensions (Å) | 49.7, 55.0, 127.6 | 41.5, 87.9, 105.6 | 71.7, 71.7, 133.2 | 71.3, 71.3, 133.5 |
| Unit Cell Angles (°) | 90, 90, 90 | 90, 90, 90 | 90, 90, 120 | 90, 90, 120 |
| Mosaic Spread (°) | 1.01 | 0.47 | 0.10 | 0.04 |
| **Reflection Data** |  |  |  |  |
| Resolution Range (Å) | 63.80-2.37<br>(2.46-2.37) | 67.57-1.75<br>(1.78-1.75) | 35.00-2.12<br>(2.26-2.12) | 35.00-1.88<br>(2.00-1.88) |
| Unique Reflections | 14847 (1507) | 39936 (2158) | 12136 (2051) | 17003 (2741) |
| *R<sub>sym</sub>* | 0.091 (0.932) | 0.147 (0.921) | 0.138 (0.991) | 0.081 (0.493) |
| I/σ(I) | 13.6 (2.0) | 9.7 (2.4) | 26.5 (6.0) | 37.0 (8.6) |
| CC½ | 0.998 (0.565) | 0.994 (0.795) | 0.999 (0.940) | 1.000 (0.986) |
| Completeness (%) | 99.9 (99.9) | 100.0 (100.0) | 100.0 (100.0) | 99.7 (98.5) |
| Multiplicity | 8.5 (6.6) | 11.8 (11.9) | 37.9 (39.7) | 38.7 (38.9) |
| Wilson B (Å<sup>2</sup>) | 45.0 | 22.1 | 42.9 | 42.8 |
| **Refinement** |  |  |  |  |
| Resolution (Å) | 63.80 (2.37) | 52.85 (1.75) | 34.61 (2.12) | 34.45 (1.88) |
| Number of Reflections | 14804 | 37878 | 11522 | 16147 |
| *R<sub>work</sub>* | 0.2366 | 0.1905 | 0.2332 | 0.2438 |
| *R<sub>free</sub>* | 0.2924 | 0.2326 | 0.2956 | 0.2881 |
| Rms (Bond Length) (Å) | 0.005 | 0.008 | 0.007 | 0.007 |
| Rms (Bond Angles) (Å) | 0.750 | 1.758 | 1.65 | 1.77 |
| **Model Composition** |  |  |  |  |
| Protein atoms | 2909 | 2895 | 1443 | 1452 |
| Waters | 24 | 134 | 57 | 78 |
| Other | / | 126 | 15 | 5 |
| **Model B-factors** |  |  |  |  |
| Protein atoms (Å<sup>2</sup>) | 60.3 | 32.2 | 44.6 | 33.3 |
| Waters (Å<sup>2</sup>) | 51.7 | 34.9 | 39.1 | 34.8 |
| Other | / | 56.4 | 62.4 | 25.1 |
| **Ramachandran Statistics** |  |  |  |  |
| Favoured (%) | 95.6 | 92.4 | 97.2 | 97.8 |
| Allowed (%) | 3.9 | 7.3 | 2.3 | 2.2 |
| Outliers (%) | 0.6 | 0.0 | 0.6 | 0 |

Values in parentheses indicate the outer resolution bin. Reflection data values are as reported by Aimless (56) for 33DL and 33DN or XDS (62) for 33DM and 33DO. Refinement statistics are as reported by Refmac (61) or Phenix (59) for 33DL. Ramachandran statistics are from Rampage (63).

**Table S2.**
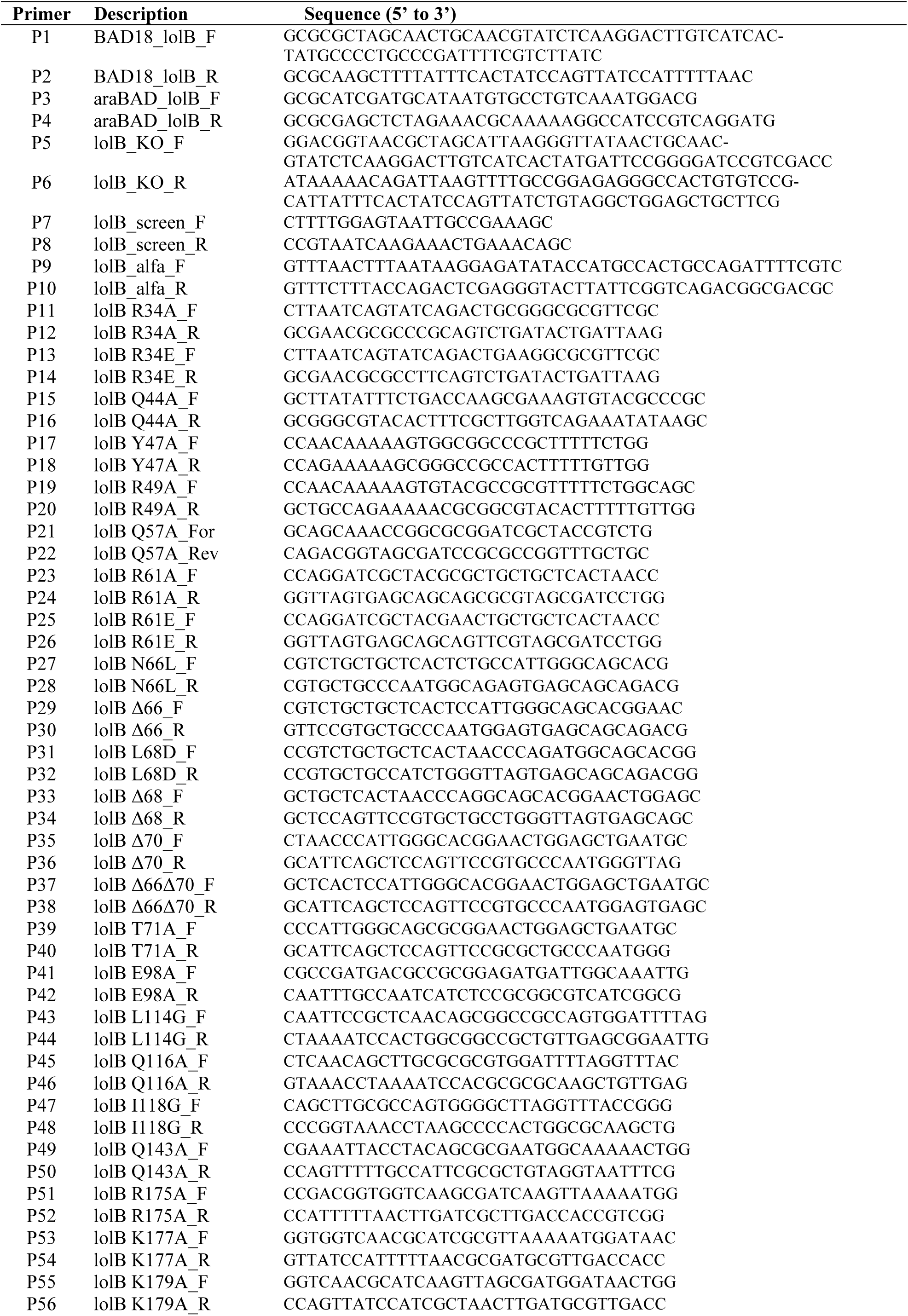
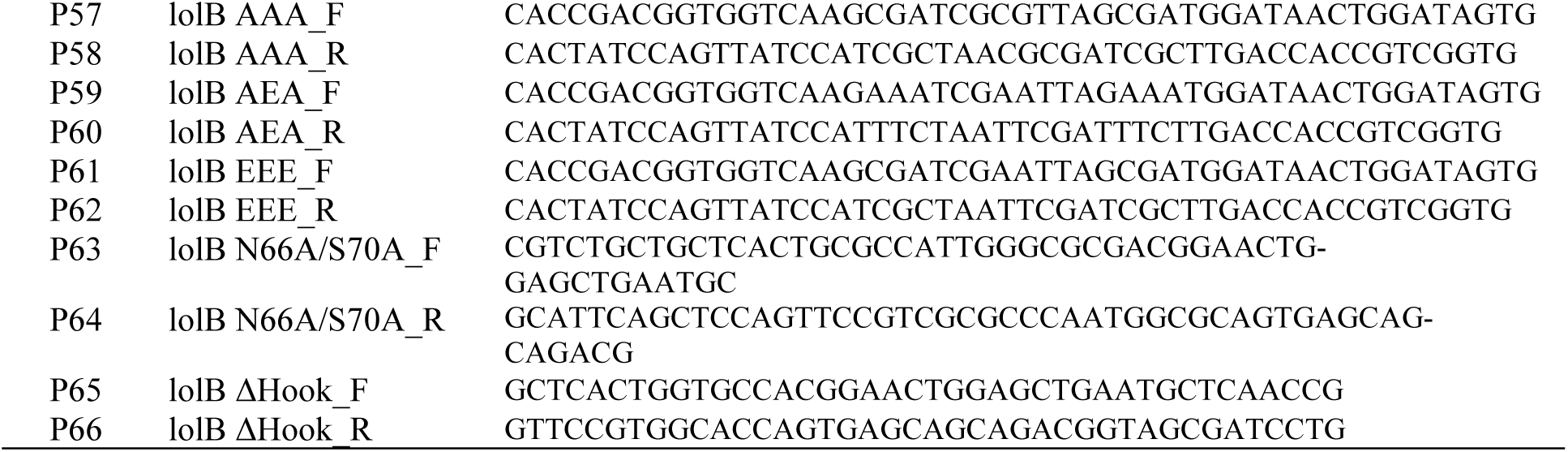
List of primers for PCR amplification.

**Table S3.** List of plasmids.

| Name | Description | Reference |
|---|---|---|
| pLDR8 | *int* gene expression vector, helper plasmid | (48) |
| pLDR9 | Cloning vector for integration into attB, kan resistant | (48) |
| pKD13 | Kan cassette template for lambda red recombination | (49) |
| pSIM5 | Expression of lambda red recombination genes | (50) |
| pCDFDuet | Expression plasmid | Novagen |
| pTH24:TEVSH | Expression of His-tagged TEV protease | (53) |
| pET28-LolB<sub>His</sub> | Expresses *E. coli* LolB (residues 23-207) with a thrombin cleavable N-terminal His-tag | (27) |
| pET28-LolB (XnY)<sub>His</sub> | Expresses *E. coli* LolB (residues 23-207) with a thrombin cleavable N-terminal His-tag, residue X at position n mutated to residue Y | This report |
| pCDF-LolB<sub>alfa</sub> | Expresses full-length LolB with a C-terminal Alfa tag | This report |
| pCDF-LolB (XnY)<sub>alfa</sub> | Expresses full-length LolB with a C-terminal Alfa tag residue X at position n mutated to residue Y | This report |
| pET28-LolB(ΔHook)<sub>His</sub> | Expresses *E. coli* LolB (residues 23-207) with a thrombin cleavable N-terminal His-tag, with residues 66-70 replaced by a GA linker | This report |
| pET28-LolB(L114G)<sub>TEV/His</sub> | Expresses *E. coli* LolB (residues 23-207) with TEV cleavage site downstream of N-terminal His-tag, and L114G mutation | This report |
| pET28-LolA | Expresses LolA (residues 22-203) with an N-terminal, thrombin cleavable His-tag | (27) |
| pET28-LolA (R43L) | Expresses LolA (residues 22-203) with an N-terminal, thrombin cleavable His-tag and R43L mutation | (27) |
| pCDF-LolA<sub>Strep</sub>-Pal<sub>WT(octa)</sub> | Co-expresses LolA with C-terminal Strep-tag (WSHPQFEK) and Pal with TEV cleavage site (ENLYFQS) upstream of a C-terminal octahistidine-tag, both separated by a GS linker | (32) |
| pCDF-LolA(R43L)<sub>His</sub>-Pal | Co-expresses LolA where residue 43 is mutated to leucine, with C-terminal His-tag and Pal with no tag | This report |
| pCDF-LolA<sub>Strep</sub>-Pal<sub>TEV/FL2</sub> | Co-expresses LolA with C-terminal Strep-tag (WSHPQFEK) and Pal with an internal TEV cleavage site at position 28 in the mature sequence and a C-terminal His-tag | (32) |

## Supplementary Movies

**Movie S1. Representative electron density for the R43L LolA - LolB structure.** LolA R43L is shown in blue and LolB in yellow. The weighted 2|*F_o_*|-|*F_c_*| electron density map is represented as a grey mesh contoured at 1 σ.

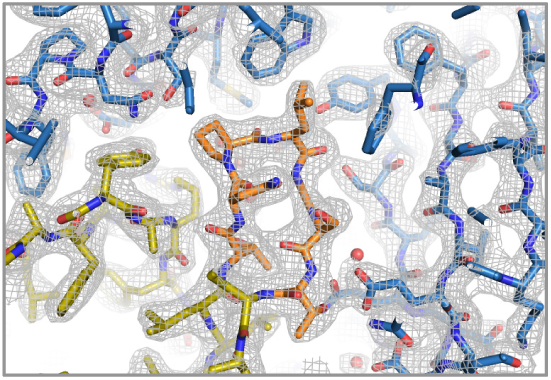

**Movie S2. Representative electron density for the LolB L114G - lipoprotein structure.** Protein residues are shown as yellow sticks and lipoprotein ligand is cyan. The crystal asymmetric unit contains two LolB molecules. The first half of the movie is focused on chain B and the second part on chain A. The weighted 2|*F_o_*|-|*F_c_*| electron density map is represented as a grey mesh contoured at 1 σ. The lipoprotein polder omit map is shown as a blue mesh contoured at 3 σ.

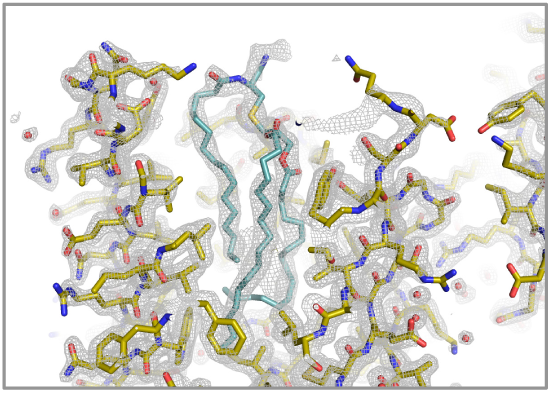

**Movie S3. Molecular morph showing the sequential transition of LolB from free to LolA-associated and lipoprotein-liganded conformations.** The movie shows LolB alternating between its unbound conformation (1IWM), bound to LolA (blue, 33DL) and liganded to a lipoprotein (cyan, 33DM). A top-down view is displayed with Lol proteins represented as cartoon (top left panel) or as surface (top right panel) showing the cavity entrance of LolB. The bottom left and right panels respectively display a side-on view of LolB and a zoom-view on the helix α3. Interactions controlling the protein conformational changes are displayed as sticks with distances ≤ 3.5 Å represented as dashed lines.

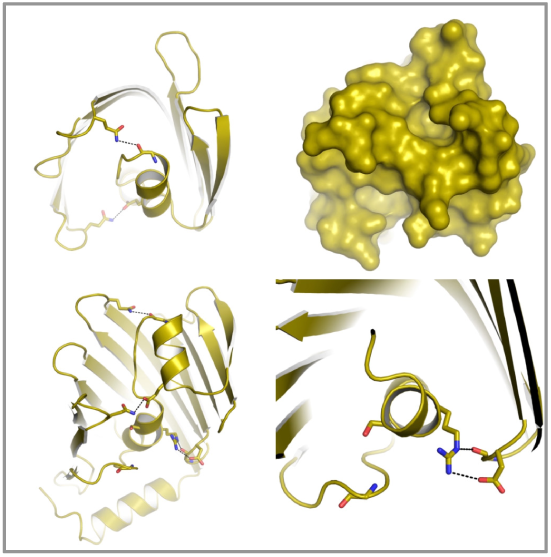

## Notes

### Competing Interest Statement

The authors have declared no competing interest.

### Summary of Updates

This version of the manuscript has been revised to update the title, minor changes to the text and Figures 4 and 5 and the introduction of two new supplementary figures.

